# Phosphorylation of serines 287/288 in DEK regulates cell-type-specific chromatin occupancy and compaction

**DOI:** 10.64898/2026.01.19.700267

**Authors:** Gongjie Wu, Matthias Meister, Sandra Reissner, Pia J. Müller, Ferdinand Kappes

## Abstract

The conserved multifunctional chromatin modulator and oncogene DEK exhibits context-dependent genomic binding and function, but how these activities are regulated in cancer remains poorly understood. Using multi-omics and biochemical approaches, we find that while DEK predominantly occupies promoter-proximal regions in HeLa cells and primary melanocytes, its chromatin binding is dramatically reduced in melanoma cell lines—despite DEK overexpression. We attributed this to CK2-mediated phosphorylation, which governs DEK chromatin association and transcriptional output in a cell-type-specific manner. Phosphoproteomics identified 34 phosphorylation sites, including S287 and S288 within the DEK C-terminal DNA-binding domain. Strikingly, CK2 inhibition and concomitant loss of phosphorylation at S287/S288 triggered DEK redistribution to promoter regions, coinciding with transcriptional repression of oncogenic pathways and global chromatin compaction. Melanoma subtypes showed divergent responses: NRAS-mutant cells displayed dynamic, phosphorylation-dependent DEK redistribution, whereas BRAF-mutant cells lacked detectable DEK binding. Our work establishes DEK as a phosphorylation-sensitive regulator of chromatin states, with CK2-mediated modification orchestrating its tumor-specific regulatory functions. These findings nominate phospho-DEK as a potential biomarker and therapeutic target in melanoma and possibly other cancers.

## Introduction

Besides histones, the large and heterogenous group of non-histone chromosomal proteins play pivotal roles in genome organization, epigenetic regulation, and coordination of transcription, DNA replication and repair ^1^. Owing to their high functional plasticity, members of this group dynamically shape chromatin structure, often via membrane-less biomolecular condensates^2^, thereby acting as context-dependent rheostats. Perturbations through mutations or post-translational modifications (PTMs) can markedly alter their activity in development and cancer.

DEK, a still enigmatic and multifunctional non-histone chromosomal protein, is an evolutionarily conserved, biochemically distinct 375-amino acid factor that has been implicated in pro-tumorigenic functions in a wide variety of human cancers^3, 4^, including melanoma ^5–7^. Unlike classical oncogenes, and safe for chromosomal translocations in leukemias ^8–11^, *DEK* is rarely mutated at the gene level, thus its oncogenicity likely arises from altered expression and dynamic, context-specific post-translational modifications ^12–16^. Distinct domains in DEK, regulated by a myriad of post-translational modifications, mediate DNA and chromatin interactions as well as its safeguarding and multitude of regulatory roles (**Fig. 1a**).

**Figure 1.**
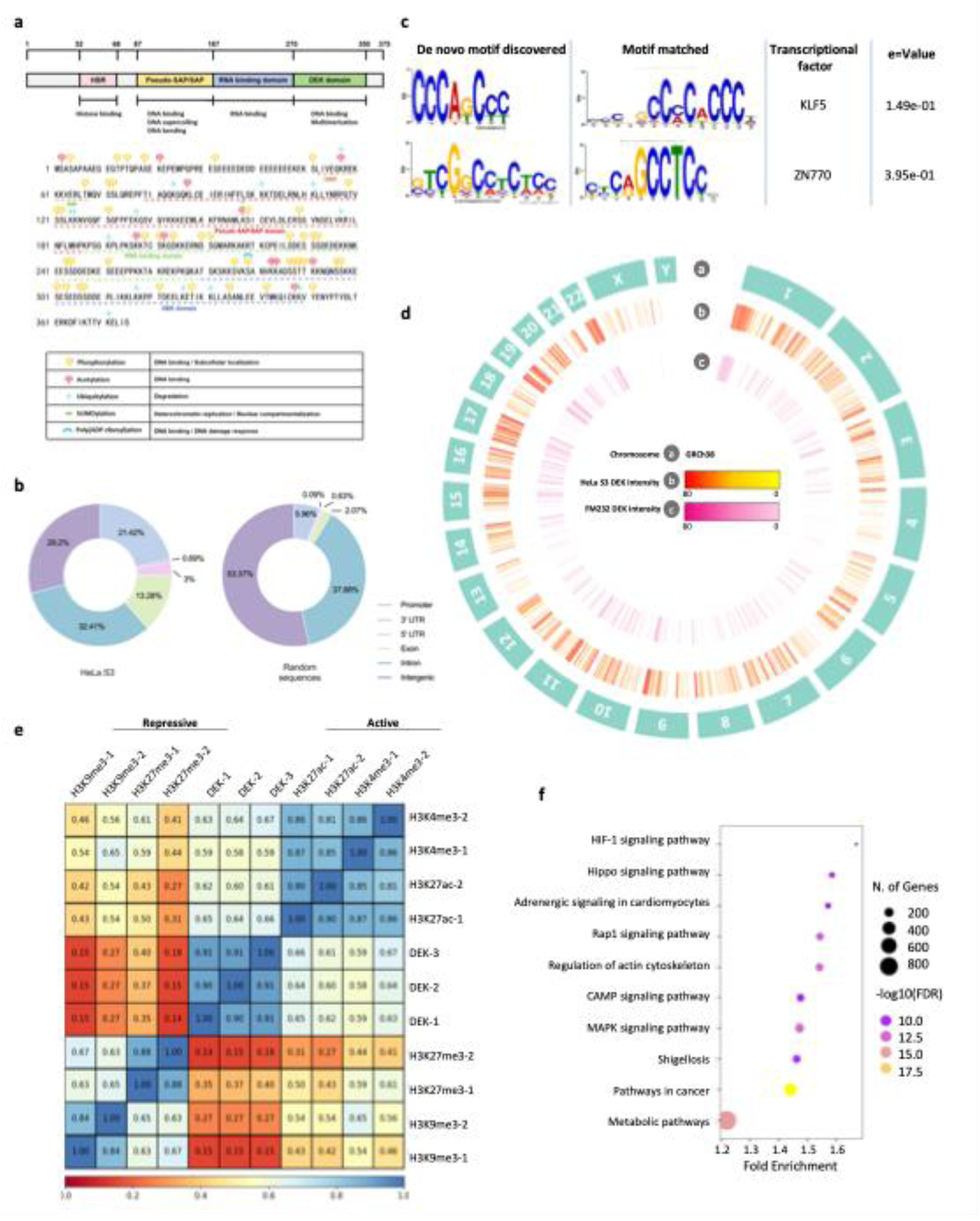
Chromatin binding landscape of DEK in HeLa S3 cells. **a,**Schematic of human DEK functional domains with annotated post-translational modification (PTM) sites (compiled from UniProt, PhosphoSitePlus, and ^14, 16, 18, 20–22, 70–72^). **b,** Genome-wide distribution of DEK ChIP-seq peaks (3 replicates) in HeLa S3 cells showing enrichment at regulatory elements (promoter-proximal and distal intergenic regions) compared with randomized genomic background. **c,** *De novo* motif analysis of the top DEK peaks identifies two enriched GC-rich motifs (motif 1: 5′-CCCAGCCC-3′, E = 1.49e-01; motif 2: 5′-GTCGGCCTCTCC-3′, E = 3.95e-01). **d,** Genome-wide distribution of merged DEK ChIP-seq peaks in HeLa S3 and FM232 (melanocytes) cells visualized by Circos; heatmap intensity indicates relative enrichment. **e,** Spearman correlation of DEK occupancy with representative histone modifications (H3K4me3, H3K9me3, H3K27me3 and H3K27ac) using ENCODE reference datasets (accession: ENCBS075PNA). **f,** KEGG pathway enrichment of DEK-associated genes in HeLa S3 cells; bubble size denotes gene counts and fold enrichment is indicated on the x axis. Pathways with FDR < 0.05 are shown.

Earlier work highlighted that DEK binding induces chromatin compaction and changes in DNA topology, thereby altering chromatin conformation and promoting chromatin compaction ^17–26^, which in part depends on its ability to bind to DNA^19^. More recently, two cryo-EM studies have solved the DEK-nucleosome structure at high resolution, revealing a unique tripartite binding mode in which the histone-binding motif (HBM) interacts with the acidic patch and other histone surfaces, the unique pseudo-SAP/SAP box binds the nucleosomal dyad and linker DNA, and a lysine-rich motif (LRM) contributes to DNA binding ^27, 28^. Together with previous data, it is now clear that DEK functions as a chromatin architectural protein, capable of promoting higher-order chromatin compaction at least under unmodified conditions.

This fundamental chromatin-shaping ability appears to be utilized in a developmental stage-and cell type specific-manner. Moreover, it appears that DEK functions may be highly versatile, as reflected by the presence of a shorter DEK2 isoform ^29^ as well as diverse post-translational modifications, including SUMOylation^18^, acetylation ^13^, poly(ADP-ribosyl)ation ^14, 16^, and phosphorylation ^22^, all of which reported to affect chromatin/DNA association of DEK as well as its secretion ^15, 16, 30, 31^. A notable example of regulation at the post-translational level is DEK’s association with hTERT, which occurs specifically in its dephosphorylated state and regulates hTERT transcriptional activity, underscoring the regulatory significance of PTMs in controlling DEK activities ^32^.

Further chromatin-centric studies connected DEK to epigenetic regulation. In mouse embryonic stem cells (mESC), Dek has been shown to safeguard H3K27me3 levels ^27^, which was also the case in *Arabidopsis thaliana*, yet with different overall outcomes^33^. In human cells, earlier chromatin studies primarily linked DEK to H3K9me3 ^17, 34^, but more recent evidence also points to its involvement in regulating H3K27me3 deposition ^28, 35^. Interestingly, DEK has also been reported to associate with lamin-associated domains (LADs) ^36^, reinforcing a potential role in organizing chromatin compartments. Yet, despite these strong connections, a recent proteomic study of chromatin readers showed that DEK in HeLa S3 cells has no clear-cut preference for any specific histone marks, thus appears not to be a typical reader of specific histone modifications or chromatin states ^37^ (**Extended data 1a**). Rather, the DNA- and nucleosome-binding capacity of DEK may be utilized and fine-tuned according to specific cellular contexts and requirements. However, how these context-dependent features, particularly the phosphorylation states of DEK, shape its chromatin functions and transcriptional outcomes in cancers remain poorly understood.

In this study, we combined multi-omics and biochemical analyses to elucidate the chromatin binding patterns of DEK in primary and cancer lines and its influence on accessibility and transcription. We then focused on the role of DEK phosphorylation states in shaping its chromatin activities. By perturbing DEK via CK2 inhibition, a kinase known to modify DEK, and targeted depletion, we demonstrate that loss of DEK phosphorylation, particularly at S278/288, drives DEK redistribution to promoter regions, an event that correlates with transcriptional repression and marked global chromatin compaction. Together, these findings establish DEK phosphorylation as a central determinant of its chromatin regulatory capacity in tumorigenesis and provide a framework for exploring DEK as a therapeutic target and as a regulator of epigenetic programs in development.

## Results

### Chromatin occupancy of DEK in HeLa cells

Our overarching goal in this study was to elucidate potential changes in the genomic occupancy of DEK in tumor versus primary cells. To do so, we started out by analyzing ChIP-seq peaks from HeLa S3 cells. DEK is predominantly bound to intragenic regions (77.27%), with a strong enrichment at promoters (21.42% of peaks within ±3 kb of TSSs; ∼4-fold enrichment over background; **Fig. 1b**), yet also to distal intergenic regions (29.2% of peaks), indicating direct and distal roles in gene regulation. Circos plots revealed genome-wide and broad chromosomal distribution, with hotspots near gene-dense regions (**Fig. 1d**). Interestingly, comparison to published DEK-ChIP-seq datasets from U937 ^38^, 786-O ^27^, HEK293 cells ^28^ and MCF7 (GEO accession GSE164429) revealed not only differences in genomic feature distribution but also striking divergence in genome-wide occupancy patterns (**Extended data Fig.1c-f**), clearly indicating pronounced cell-type and context specific functions of DEK in gene regulation. Motif analysis of highly enriched DEK binding sites identified GC-rich sequences (CCCAGCCC and GTCGGCCTCTCC; e-value= 1.49e-01 and 3.95e-01; **Fig. 1c**), resembling binding motifs for transcription factors KLF5 and ZNF770. This contrasts with reported AT-rich preferences observed *in vitro*, suggesting, again, context-dependent chromatin association of DEK ^20, 24^. Co-occurrence analysis of ChIP-seq peaks demonstrated that DEK overlaps with active marks (H3K4me3, H3K27ac) but minimally with repressive H3K9me3 or H3K27me3 modifications (**Fig. 1e**). KEGG pathway enrichment analysis linked DEK-bound genes to Hippo, Rap1 and MAPK signaling (*FDR*< 0.05; **Fig. 1f**), underscoring its potential oncogenic roles.

### DEK loss remodels chromatin accessibility and transcriptional programs

Next, we applied a multi-omics approach combining ATAC-seq (Assay for Transposase-Accessible Chromatin using sequencing), gene expression microarrays (Affymetrix HTA 2.0) from both wild-type (WT) and DEK full-length depleted (1E7, expressing truncated DEK, **Extended data Fig. 2**) HeLa S3 cells with our DEK-ChIP-seq data from WT cells. Principal component analysis (PCA) of microarray data revealed distinct clustering between WT and 1E7 cells (PC1: 74.5% variance; **Fig. 2b**), with 1,339 differentially expressed genes (DEGs; 819 up, 520 down; FDR < 0.05; **Fig. 2a**). Enrichment analysis revealed upregulation of PI3K-AKT and ECM pathways, while downregulated genes were associated with cardiomyopathy-related pathways (**Fig. 2c, d**). ATAC-seq on these cells (two biological replicates per condition) revealed a global increase in TSS-proximal chromatin accessibility upon loss of full-length DEK (**Fig. 2e**), suggesting DEK normally restricts access to such regulatory elements. Unsupervised clustering of ATAC-seq signals (±5 kb from TSS) identified five distinct chromatin states. Two clusters exhibited DEK-dependent accessibility: cluster 1 gained accessibility, while cluster 2 lost accessibility upon depletion of full-length DEK (**Fig. 2f**). This bimodal response indicates DEK functions as both a chromatin opener and repressor in a locus-specific manner, potentially mediated by distinct DEK pools. The remaining clusters displayed stable accessibility, implying independence from DEK-mediated regulation. Integration of ATAC-seq and microarray datasets uncovered 134 genes showing concurrent increases in both chromatin accessibility (log2 fold change > 1, FDR < 0.05) and expression (log2 fold change > 0.5, FDR < 0.05), alongside 63 genes exhibiting coordinated decreases (**Fig. 2g**). Notably, the up-regulated gene set showed significant enrichment (FDR < 0.01) in melanomagenesis-related pathways (KEGG pathway hsa04916), including key regulators such as PLCB1 (phospholipase C beta 1, log2FC = 1.2), CAMK2D (calcium/calmodulin-dependent protein kinase II delta, log2FC = 1.1), and CREB3L2 (cAMP responsive element binding protein 3 like 2, log2FC = 0.9; **Fig. 2i**). These findings are particularly relevant given the established role of DEK as an oncogene in melanoma, suggesting it may regulate melanomagenesis programs through direct control of chromatin accessibility and gene expression.

**Figure 2.**
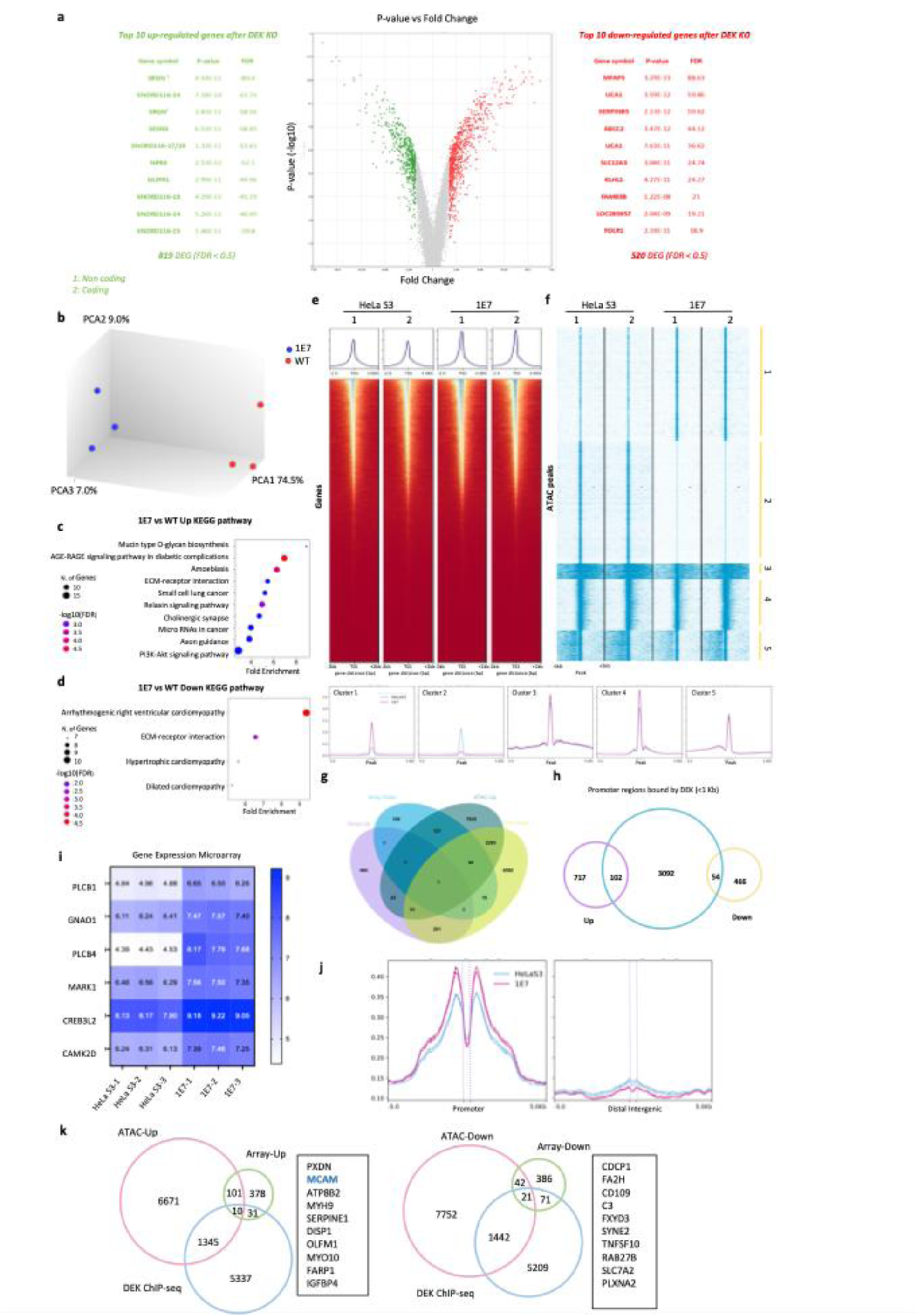
DEK loss reshapes transcriptional programs and chromatin accessibility in HeLa S3 cells. **a,** Volcano plot of differentially expressed genes (DEGs) from expression microarray profiling (3 replicates) comparing wild-type (WT) and DEK-depleted (1E7) HeLa S3 cells (FDR < 0.05); upregulated genes are shown in red and downregulated genes in green. **b,** Principal component analysis (PCA) of expression microarray profiles from WT and 1E7 cells showing separation along PC1. KEGG pathway enrichment of genes upregulated (**c**) or downregulated (**d**) upon DEK depletion; top pathways with FDR < 0.05 are shown. **e,** Genome-wide ATAC-seq signal displayed as a heatmap for WT and DEK-depleted cells (duplicates), with the average profile around transcription start sites (TSS) shown above. **f,** Unsupervised clustering of ATAC peaks identifies five accessibility patterns; average accessibility profiles for each cluster are shown below. **g,** Overlap between DEGs inferred from ATAC-seq-associated genes and DEGs from expression profiling; bar plot summarizes up- and downregulated categories. **h,** Overlap between promoter-proximal DEK binding (±1 kb from TSS) and DEGs following DEK depletion. Numbers indicate DEGs with promoter DEK occupancy (upregulated: 102; downregulated: 54) versus without promoter occupancy (upregulated: 717; downregulated: 466). **i.** Heatmap of expression microarray signals for genes in the KEGG melanogenesis pathway in WT and DEK-depleted (1E7) HeLa S3 cells; color intensity reflects relative expression (blue indicates higher expression). **j,** Aggregate ATAC–seq profiles showing accessibility changes after DEK depletion at promoter regions (top) and distal intergenic regions (bottom). **k,** Integrated analysis of DEK ChIP-seq, ATAC-seq and expression profiling identifies genes with concordant changes in accessibility and transcript levels upon DEK depletion; the 10 genes in the three-way overlap (DEK occupancy, co-upregulated accessibility and co-upregulated expression) are listed.

### Context-dependent regulation of chromatin accessibility and transcription by DEK

Further examination of binding patterns of DEK revealed 6,615 promoter-proximal sites (within 1 kb of transcription start sites), with 156 of these associated with genes differentially expressed following depletion of full-length DEK. Among these DEK-bound targets, up-regulated genes (102 genes, 65.4%) predominated over down regulated genes (54 genes, 34.6%), supporting a primary role for DEK as a transcriptional repressor in this context (**Fig. 2h**). Meta profile analysis of ATAC-seq signals centered on DEK-bound regions revealed striking context-dependent roles of DEK in shaping chromatin accessibility. At promoter-proximal regions (≤3 kb from TSS), DEK full-length depletion led to reduced accessibility precisely at the DEK-occupied core (average tag density increase of 8%), but increased accessibility in flanking regions (average tag density reduction of 22.9%), indicating that DEK normally promotes focal chromatin accessibility at its binding sites while constraining chromatin opening in the surrounding promoter context (**Fig. 2j**). In contrast, distal intergenic peaks exhibited a uniform loss of accessibility in 1E7 cells (average tag density reduction of 13%), suggesting that DEK may support open chromatin configurations at distal regulatory elements (**Fig. 2j**).

Building on this, a three-way integration analysis identified two distinct classes of DEK-regulated genes: *(i)* 10 loci that lost DEK binding, gained increased chromatin accessibility (ATAC-seq log2FC > 1), and showed transcriptional upregulation (microarray log2FC > 0.5), and *(ii)* 21 loci that exhibited reduced accessibility accompanied by transcriptional downregulation upon DEK loss. Notably, amongst the upregulated group, MCAM (Melanoma Cell Adhesion Molecule, CD146, log2FC = 1.8) stood out as a well-characterized mediator of melanoma progression and metastasis (**Fig. 2k**). Together, these results indicate that DEK exerts dual, context-specific roles in chromatin regulation - repressing transcription at promoters while maintaining accessibility at distal sites - and that its disruption rewires oncogenic programs, including activation of melanoma drivers such as MCAM. The context-dependent nature of DEK’s activity - acting as both a repressor and activator depending on genomic location - highlights its importance as a modulator of chromatin dynamics and suggests it may serve as a critical node in the maintenance of oncogenic gene expression networks.

### Cell-type-specific DEK occupancy distinguishes normal and malignant contexts

DEK is overexpressed in invasive melanoma ^5, 39^ and promotes tumorigenesis by enhancing proliferation and chemoresistance ^6, 7^. To now assess its role in melanoma progression, we initially compared DEK chromatin occupancy between primary melanocytes (FM232) and HeLa S3 cells using ChIP-seq. In both cell types, DEK predominantly localized to promoter regions (**Fig. 3a** and **1d**), correlating with active histone marks (H3K4me3, H3K27ac; **Fig. 3b**). Differential binding analysis revealed 15,405 DEK-bound regions with significant cell-type-specific differences (FDR < 0.01), with FM232 exhibiting fewer peaks than HeLa S3 (**Fig. 3c**). GO enrichment analysis indicated that DEK-bound genes in FM232 were associated with transcriptional regulation and chromatin organization, whereas in HeLa S3, they were linked to transmembrane transport and cell signaling pathways (**Fig. 3d**). These findings suggest that even though DEK maintains conserved promoter-proximal binding patterns, its target gene network diverges substantially in normal versus malignant cells, which may contribute to oncogenic reprogramming.

**Figure 3.**
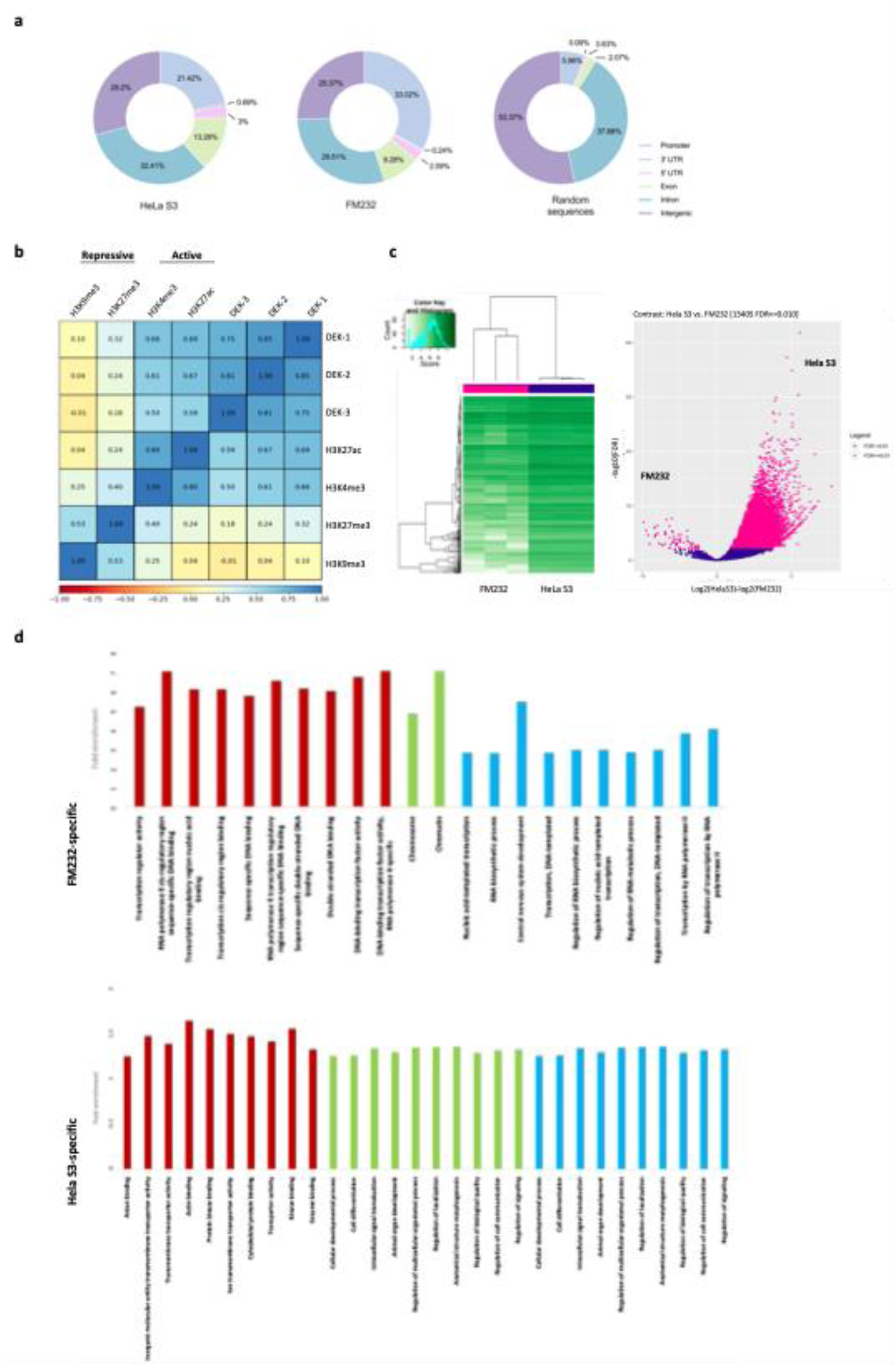
DEK occupancy differs between FM232 and HeLa S3 cells and is associated with distinct chromatin contexts. **a,** Genomic annotation of DEK ChIP-seq peaks in FM232 and HeLa S3 cells (GRCh38/hg38), compared with randomized genomic background. **b,** Spearman correlation of DEK occupancy with representative histone modifications (H3K4me3, H3K9me3, H3K27me3 and H3K27ac) in FM232 cells using ENCODE reference datasets (accessions ENCBS075PNA and ENCBS041UOV). c, Differential DEK binding between HeLa S3 and FM232 cells. Volcano plot shows log2 fold change in DEK enrichment versus statistical significance (−log10 FDR); significantly differential regions (FDR < 0.01) are highlighted. d, Gene ontology (GO) enrichment of genes associated with cell-type-specific DEK-bound regions. GO terms are shown for FM232-specific (top) and HeLa S3-specific (bottom) DEK-bound genes; fold enrichment is indicated.

### Dramatically altered chromatin association of DEK in melanoma cells

To finally elucidate a potential role for DEK in melanomagenesis, we performed ChIP-seq in five melanoma cell lines (SK-Mel-19, SK-Mel-29, SK-Mel-103, UACC-62, UACC-257, see **Extended data 3a** for details). Surprisingly, peak calling with standard thresholds (q<0.05) yielded essentially no reproducible peaks, regardless of p53 status or key mutations in signaling molecules in these melanoma cell lines as compared to primary melanocytes (FM232) and HeLa S3 cells (**Fig. 4a**). Control experiments excluded technical issues, as DEK antibodies efficiently immunoprecipitated endogenous protein from these cells (**Extended data Fig.3b**). Given this remarkably peculiar result, we first broadly analyzed the chromatin association of DEK as defined by micrococcal nuclease (MNase) accessibility using either time course experiments or a differential extraction method^40^, which we first established in HeLa S3 cells (**Extended data Fig. 3c**). Whereas DEK predominantly remained in the immobilized chromatin fraction in primary melanocytes (**Fig. 4b, Extended data Fig. 4a**), almost all DEK was found in the mobilized chromatin fractions in SK-MEL-103 (**Fig. 4b**), with other cells lines showing intermediate patterns (**Fig. 4c; Extended data Fig. 4a**). Remarkably, HeLa S3 cells displayed a melanoma-like extraction pattern yet retained abundant ChIP peaks, suggesting pronounced qualitative differences in the DEK populations within these cells.

**Figure 4.**
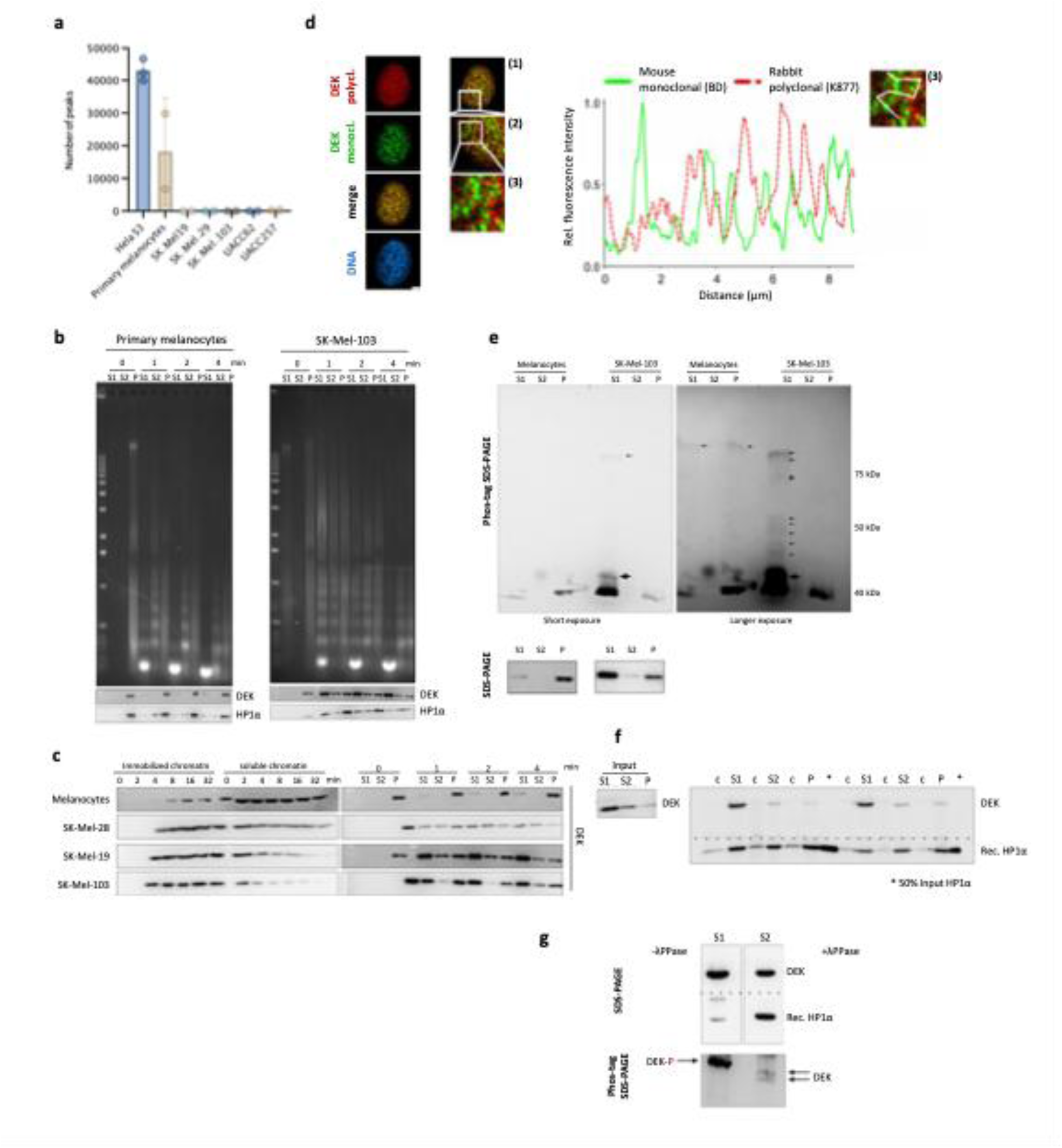
Melanoma cells exhibit reduced chromatin-bound DEK and increased DEK phosphorylation associated with impaired HP1α binding. **a,** DEK ChIP-seq peak numbers in HeLa S3 cells, primary melanocytes and melanoma cell lines (SK-Mel-19, SK-Mel-29, SK-Mel-103, UACC-62 and UACC-257; 3 replicates for each cell line). **b,c,** MNase accessibility–based chromatin fractionation and differential extraction to assess DEK chromatin association in primary melanocytes and melanoma cells. **d,** BJ5ta fibroblasts were immunolabeled with DEK-specific antibodies. Confocal image of an interphase nucleus labeled with a mouse phosphorylation-sensitive monoclonal α-DEK antibody (green) and rabbit polyclonal α-DEK antibody (red). Scale bar: 5 μm. Zoom-in view of inset of merged image shown on the left. right panel: fluorescence intensity profiles of DEK-specific signal (green: mouse monoclonal; red-dashed: rabbit polyclonal) along a line as shown. **e,** Phos-tag SDS–PAGE of endogenous DEK found in S1, S2 and P chromatin fractions in primary melanocytes and SK-Mel-103 cells showing increased DEK phosphorylation in SK-Mel-103. The bottom panel shows the same samples resolved by standard SDS-PAGE, **f,** HP1α binding assay. DEK from indicated fractions from SK-MEL103 cells was immunoprecipitated DEK and incubated with recombinant HP1α. C indicates beads only. Samples were analyzed by immunoblotting. Input is shown on the left. g, HP1α binding assay as in (f) with DEK from the S1 fraction treated ±λ protein phosphatase (λPPase) prior to HP1α binding.

### CK2-driven phosphorylation governs chromatin association and functional partitioning of DEK

The domain organization and PTM distribution of DEK are summarized in **Figure 1a**. Interestingly, probing total cell lysates with a monoclonal DEK antibody, known to preferentially recognize phosphorylated DEK species ^16^, showed increased recognition of DEK in SK-Mel-103 along with generally increased CK2 expression in the chosen melanoma cell lines (**Extended data Fig. 4b**), consistent with previous reports ^41^. This prompted us to investigate the phosphorylation status of DEK in melanocytes versus SK-MEL-103. Phos-tag SDS-PAGE revealed markedly increased phosphorylation of DEK in SK-Mel-103, particularly within the active chromatin-associated S1 fraction (**Fig. 4e**). Mass spectrometry of immunoprecipitated DEK from nuclear lysates confirmed extensive phosphorylation, with 34 sites in SK-Mel-103 and 18 in HeLa S3 (MS data could not be obtained from primary melanocytes) with 12 overlapping sites, 3 exclusive sites (131S, 287S, 288S in SK-Mel-103), and 3 sites (S32, S251, S307) as known CK2 target sites (**Fig. 5h**). CK2 is a well-established DEK-interacting kinase and major regulator of its phosphorylation ^42, 43^. CK2 orchestrates DEK’s histone chaperone activity enabling H3.3 deposition and nuclear receptor coactivation at euchromatic loci, whereas dephosphorylated DEK supports heterochromatin integrity via interactions with heterochromatin protein 1α (HP1α) ^17, 44^. This dual regulation positions DEK as a dynamic task-specific chromatin modulator, with its post-translational status dictating spatial and functional partitioning - a mechanism potentially disrupted or altered in tumorigenesis.

**Figure 5.**
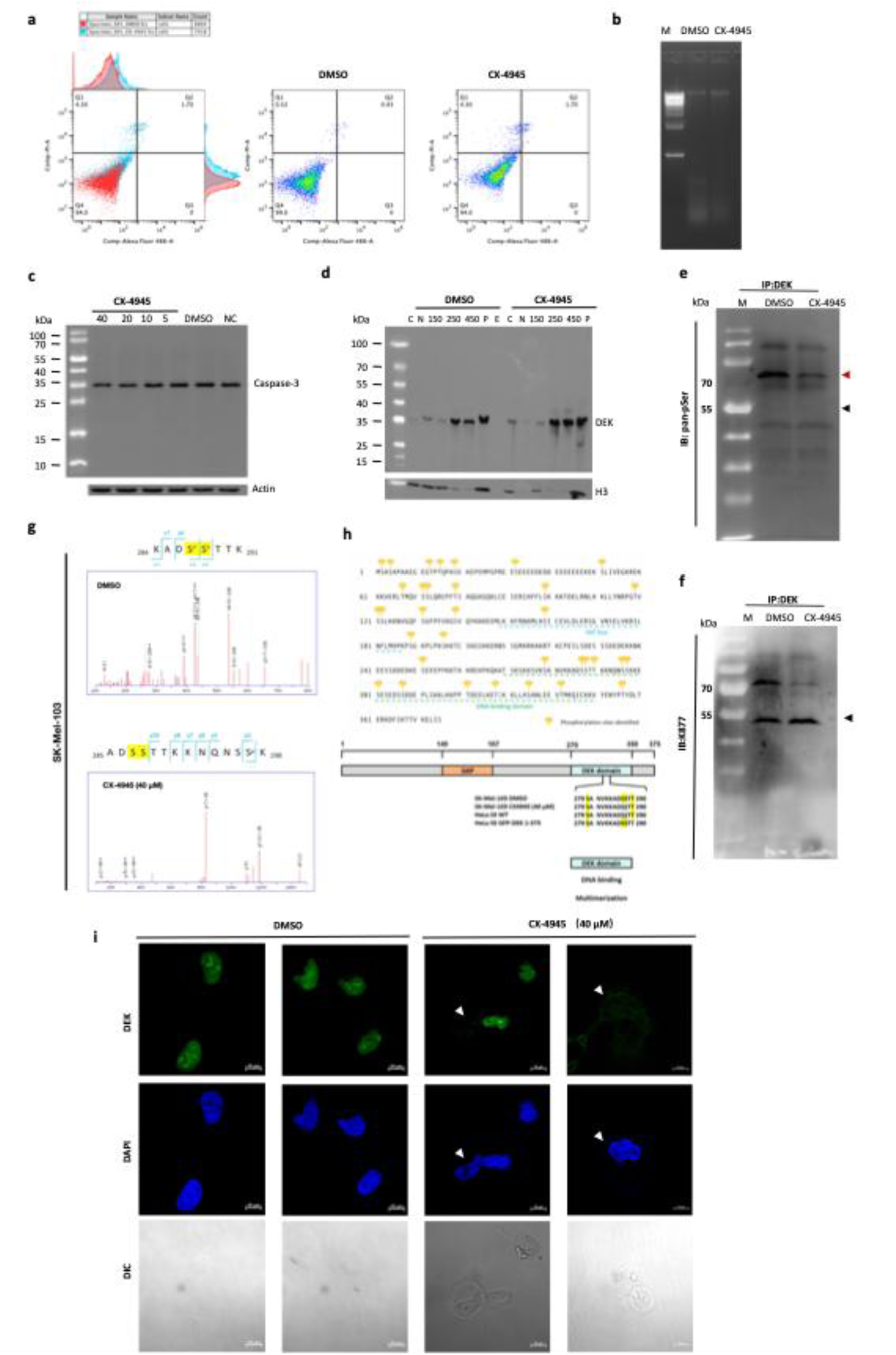
CK2 inhibition modulates DEK phosphorylation and subcellular distribution in SK-Mel-103 cells. **a,** Annexin V–FITC/propidium iodide (PI) apoptosis assay of SK-Mel-103 cells treated with CX-4945 (40 µM, 24 h) or DMSO. Scatterplots depict four different quadrant: viable cells (Annexin V⁻/PI⁻, Q4), early apoptotic (Annexin V⁺/PI⁻, Q3), late apoptotic/necrotic (Annexin V⁺/PI⁺, Q2), and necrotic (Annexin V⁻/PI⁺, Q1). **b,** Apoptosis DNA fragmentation analysis in SK-Mel-103 following CX-4945 treatment. **c,** Immunoblot analysis of caspase-3 cleavage in SK-Mel-103 cells treated with CX-4945 (5–40 µM) or DMSO; actin serves as loading control. **d,** Subcellular fractionation of SK-Mel-103 cells into cytosolic, nucleosolic and salt-extracted chromatin fractions (100, 250 and 450 mM NaCl), followed by immunoblotting for DEK. **e,** Immunoprecipitation of endogenous DEK from SK-Mel-103 cells treated with CX-4945 (40 µM, 24 h) or DMSO, followed by immunoblotting with a pan-phosphoserine antibody. **f,** Re-probing of the immunoprecipitated DEK membrane with a polyclonal anti-DEK antibody (K877). Clean-Blot IP Detection Reagent was used to minimize interference from immunoglobulin heavy and light chains. **g,** Representative LC–MS/MS spectra of DEK-derived phosphopeptides from SK-Mel-103 cells treated with DMSO or CX-4945. **h,** Summary of DEK phosphorylation sites identified by LC-MS/MS and schematic of DEK domains (top); site alignment is shown for endogenous DEK from SK-Mel-103 and HeLa S3 cells, and GFP–DEK expressed in HeLa S3 cells (bottom). **i,** Immunofluorescence of SK-Mel-103 cells after CK2 inhibition (CX-4945, 40 µM, 24 h) stained for DEK (scale bars:10µm).

Indeed, DEK colocalizes with HP1α in primary melanocytes, yet to a lesser extent in melanoma cells, where colocalization with H3.3 predominated (**Extended data Fig. 5a and b**). Whereas weakly phosphorylated DEK immunoprecipitated from inactive P chromatin fractions interacted strongly with recombinant HP1α, DEK immunoprecipitated from S1 fractions showed only weak affinity (**Fig. 4f**), which was restored upon dephosphorylation with λ-phosphatase (**Fig. 4g**), confirming phosphorylation as a key regulator of DEK’s chromatin-associated functions. Even though two recent cryo-EM studies excluded C-terminal DEK regions as crucial for nucleosome binding and subsequent chromatin compaction, phosphorylation events targeting full-length DEK suppresses its DNA and chromatin binding and fosters multimerization of DEK ^21, 22, 42^, which we re-confirmed by MNase digestions of SV40 minichromosomes, sucrose density gradient analyses and native gel electrophoresis (**Extended Data Figs. 6a, b, c**). Analysis of fibroblasts with the monoclonal DEK antibody mentioned above and a polyclonal DEK antibody, revealed co-existence of clearly discernable phosphorylated and unphosphorylated DEK populations, suggesting a cell-type-specific balance of differentially post-translationally modified DEK species (**Fig. 4d**). These findings, together with the reduced ChIPability of DEK in melanoma cells, support a model in which CK2-mediated phosphorylation promotes DEK eviction from chromatin, as proposed before ^22^.

### CK2-dependent phosphorylation of DEK in melanoma cells

To test whether CK2 activity indeed drives DEK hyperphosphosphorylation in melanoma cells, we treated SK-Mel-103 cells with the ATP-competitive CK2 inhibitor CX-4945 (Silmitasertib, 40 µM, 24 h). This treatment resulted in a modest increase in apoptotic cells (**Fig. 5a**; Annexin V⁺/PI⁺, Q4 (viable) vs Annexin V⁻/PI⁻, Q2 (apoptotic/necrotic)), without detectable apoptotic DNA laddering (**Fig. 5b**) or caspase 3 cleavage (**Fig. 5c**). Pan-phosphoserine immunoblotting of immunoprecipitated endogenous DEK revealed a 70-80 kDa DEK species in control cells (DMSO), reduced by ∼48% upon CK2 inhibition (**Fig.5d, left**). Notably, conventional DEK antibodies failed to detect these species, suggesting phosphorylation-dependent epitope masking. A predominant 55 kDa DEK isoform, representing a less phosphorylated form, remained unchanged (**Fig. 5e and f**). Residual phosphorylation (∼52%) reflects incomplete CK2 inhibition or, more likely, a range of compensatory post-translational modifications by other kinases or enzymes, as observed before ^18^.

### LC-MS/MS identifies CK2-regulated DEK phosphosites

Mass spectrometry extended these findings by mapping 34 phosphorylation sites on endogenous DEK from SK-Mel-103 and HeLa S3 cells, including six novel sites (S2, T79, S139, S189, S196, T356) (**Extended data Figs. 7a, b, c**). Among these, CK2 inhibition specifically abolished phosphorylation at S287/S288 within DEK’s DNA-binding domain in SK-Mel-103 cells (**Fig. 5g and h**). Intriguingly, these sites were unmodified in endogenous DEK in HeLa S3 cells but phosphorylated in lentiviral overexpressed GFP-tagged DEK, even when a C-terminal large-scale serine-to-alanine mutant (S-11-A mutant: S-227, 230, 231, 232, 243, 244, 251, 301, 303, 306, 307-A) was expressed, suggesting rapid cell-type specific compensatory modification of ectopic protein. Functionally, overexpression of GFP-tagged DEK in SK-Mel-103 cells resulted in an initially dramatic, yet highly transient, compaction of chromatin (**Extended data Fig. 6e**), reminiscent of the drastic compaction of bacterial nucleoids upon human DEK expression, which also results in suppression of bacterial growth ^45^. This, however, was not observed for S-287/S288 to alanine or glutamic acid mutans when subjected to a bacterial growth inhibition screen (**Extended data Fig. 6d**), suggesting that these two amino acids affect overall confirmation of full-length DEK. In further support of rapid compensatory modifications, subsequent MNase chromatin fractionation at 12 hours revealed that both GFP-DEK and the S-11-A mutant localized predominantly to the mobilized chromatin fraction with little impact on global compaction, recapitulating endogenous DEK (**Extended data Figs. 6f, g**). These findings suggest that ectopically expressed DEK is rapidly modified post-translationally, including potential compensatory events as previously observed upon mutation of SUMOylation sites, thereby constraining its chromatin-compacting activity ^12^.

### CK2 inhibition alters DEK localization

To elucidate the impact of CK2 inhibition on the subcellular localization of DEK, we conducted cell fractionation assays in SK-Mel-103 cells treated with CX-4945. CK2 inhibition led to increased presence of cytosolic DEK alongside enhanced association with chromatin fractions extractable at high salt concentrations (450 mM NaCl), suggesting CX-4945-induced dephosphorylation increases DEK’s chromatin affinity while simultaneously promoting compensatory modifications that facilitate dynamic nucleocytoplasmic shuttling, potentially linked to its secretion (**Fig. 5d, Extended Fig. 9a**). Consistent with this observation, immunofluorescence imaging revealed a loss of nuclear DEK in subsets of cells, supporting the role of CK2 in mediating DEK secretion and function as a pro-inflammatory molecule (**Fig. 5i**). Together with LC-MS/MS results, these findings suggest that S287/S288 may represent key CK2-regulated phosphosites involved in modulating DEK’s chromatin interactions and dynamic subcellular distribution.

### CK2 inhibition reshapes DEK’s chromatin landscape

Building on the observation that CK2 inhibition modulates DEK phosphorylation and alters its chromatin affinity, ChIP-seq profiling in SK-Mel-103 cells further revealed that CX-4945 treatment induced a stark redistribution of DEK chromatin occupancy, with PCA analysis showing clear separation from DMSO controls (**Fig. 6a**). Although untreated SK-Mel-103 cells exhibited limited DEK binding peaks (**Fig. 4a**), DMSO exposure markedly increased the number of detectable peaks, likely reflecting the effect of DMSO on protein solvation and phase separation^46^ that may unspecifically enhance DEK’s chromatin association (**Fig. 6a**). Importantly, CX-4945-specific DEK occupancy was predominantly enriched at promoters of genes such as UFD1 and MARK3 (**Fig. 6b**), which was further validated by ChIP-qPCR (**Fig. 6d**). Extending this regimen to UACC-62 cells revealed striking subtype-specific differences: whereas aggressive NRAS Q61R-mutant SK-Mel-103 melanoma cells exhibit dynamic, phosphorylation-dependent DEK chromatin redistribution, non-aggressive BRAF V600E-mutant UACC-62 cells show no detectable DEK binding under either basal or CX-4945-treated conditions in both genome-wide ChIP-seq and targeted ChIP-qPCR analyses (<0.5 percentage input) (**Extended data Fig. 9b**). This fundamental dichotomy between melanoma subtypes likely reflects how a specific oncogenic mutation background dictates DEK’s chromatin engagement through post-translational regulation. Furthermore, cell-type dependency extended to non-transformed cells, with mammary epithelial cells displaying extensive DEK binding (∼1000 peaks) compared to minimal association in dermal fibroblasts, underscoring a lineage-specific pattern of DEK-chromatin interactions (**Extended data Fig. 9d**). Together, DEK’s chromatin association emerges as a highly context-sensitive process, rewired by lineage identity, oncogenic mutations, phase-separation sensitivity converge, with phosphorylation acting as a molecular switch that redefines its chromatin interactions in aggressive melanoma.

**Figure 6.**
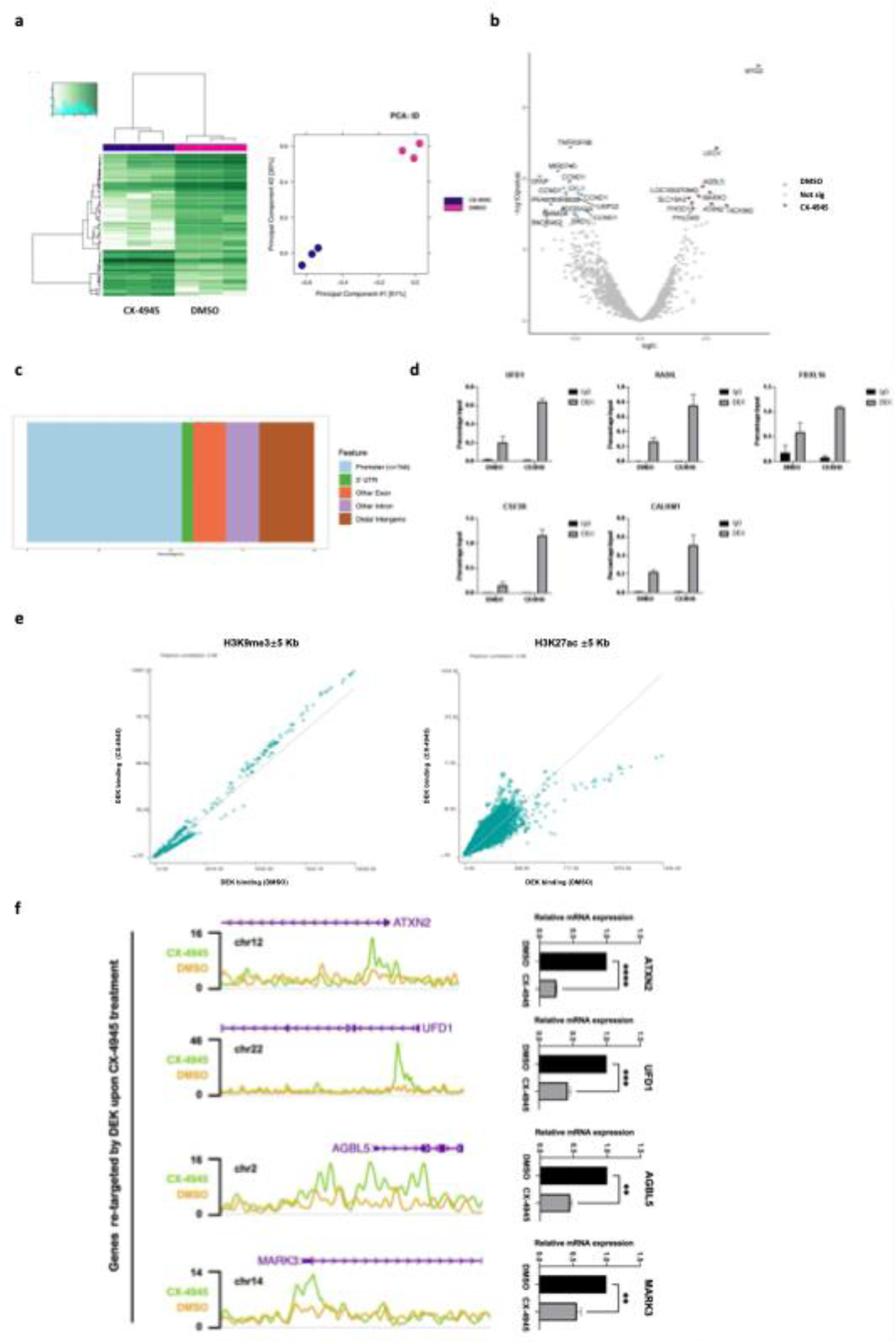
CK2 inhibition reshapes the DEK chromatin-binding landscape in SK-Mel-103 cells. **a,** DEK ChIP-seq profiling in SK-Mel-103 cells treated with DMSO or CX-4945. Heatmap and principal component analysis (PCA) summarize DEK occupancy across samples (PC1 = 61%, PC2 = 30%). **b,** Differential DEK binding analysis between CX-4945 and DMSO conditions shown as a volcano plot (log2 fold change versus −log10 FDR); significantly differential regions are highlighted. **c,** Genomic annotation of CX-4945-specific DEK-bound regions. **d,** ChIP-qPCR validation of CX-4945–specific DEK occupancy at indicated loci (UFD1, RADIL, FBXL16, CSF3R and CALHM1) using DEK ChIP and IgG control; data are shown as percent input (mean ± s.d., n = 3). **e,** Association of DEK occupancy with active (H3K27ac) and repressive (H3K9me3) chromatin features. DEK ChIP-seq tags from DMSO- and CX-4945-reated cells were counted within ± 5kb windows centered on H3K27ac or H3K9me3 peak coordinates. **f,** Representative CX-4945-associated DEK peaks in ChIP-seq tracks (left). RT-qPCR of the corresponding genes showing reduced mRNA levels upon CX-4945 treatment (right; mean ± s.d., n = 3; two-tailed Student’s t-test, *P* < 0.05, \*\**P* < 0.005).

### Phosphorylation-dependent redistribution of DEK towards repressive transcription

To further dissect the functional consequences of CX-4945-induced DEK redistribution, we compared overall DEK binding alterations in DMSO- and CX-4945-treated SK-Mel-103 cells to our own generated H3K9me3 and H3K27ac-specific CHIP-seq peaks. We found that DEK occupancy progressively shifted from the active enhancer mark H3K27ac and toward repressive marks H3K9me3 following CK2 inhibition (**Fig. 6e**), suggesting that dephosphorylation directs DEK to transcriptionally silent chromatin. Consistent with this redistribution, we found that over 50% of CX-4945-unique DEK accessible regions were localized to promoter regions (**Fig. 6c**). ChIP-seq tracks highlighted prominent enrichment of DEK at these loci, and transcriptional profiling confirmed that the corresponding genes underwent marked downregulation (**Fig. 6f**). Importantly, this downregulation was not as pronounced in DEK full-length-depleted HeLa S3 cells (1E7), underscoring a direct role for DEK in mediating CK2 inhibition-driven transcriptional silencing (**Fig. 7a and b, Extended data Fig. 8a**).

**Figure 7.**
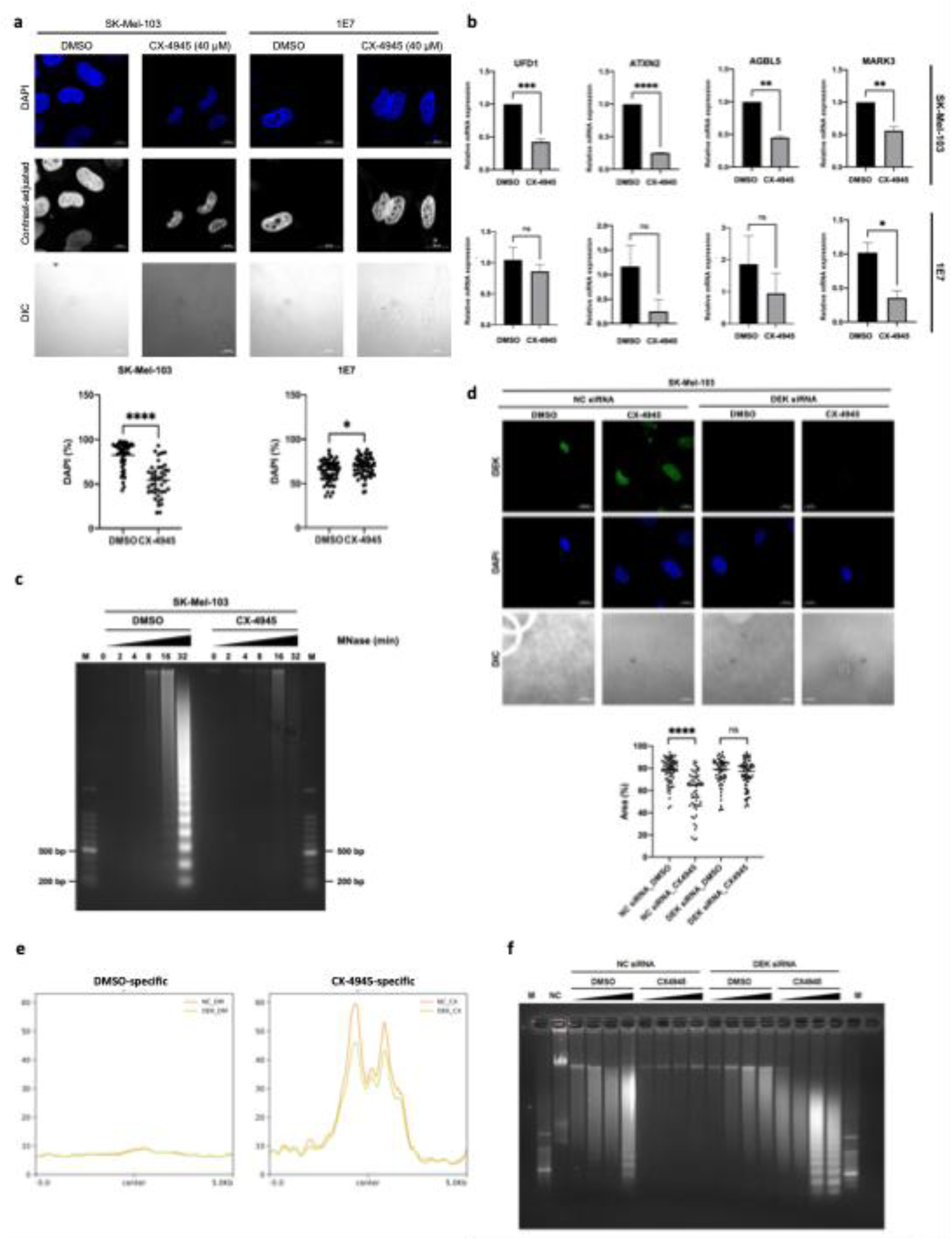
CK2 inhibition promotes chromatin compaction in SK-Mel-103 cells in a DEK-dependent manner. **a,** Confocal imaging of nuclear chromatin morphology (DAPI) in SK-Mel-103 cells and DEK-depleted 1E7 cells following CX-4945 treatment; quantification of DAPI-positive nuclear area is shown (each dot represents one nucleus; n = 43, 50, 59, 60 as indicated; two-tailed Student’s t-test, *P* < 0.05, \*\**P* < 0.005). Contrast-adjusted grayscale renderings of the DAPI channel are shown to aid visualization of chromatin compaction. Scale bar, 10 μm. **b,** RT-qPCR analysis of selected transcripts in DEK-depleted 1E7 cells versus controls following CX-4945 treatment (mean ± s.d., n = 3; two-tailed Student’s t-test, *P* < 0.05, \*\**P* < 0.005). **c,** MNase digestion time course of nuclei isolated from DMSO-or CX-4945-treated SK-Mel-103 cells; purified DNA was resolved by agarose gel electrophoresis. **d,** Immunofluorescence staining for DEK in SK-Mel-103 cells following siRNA-mediated DEK knockdown and CX-4945 treatment, with DAPI counterstaining. Scale bar, 10 μm. **e,** ATAC-seq Metaprofiles centered on DMSO-specific (left) and CX-4945-specific (right) DEK binding sites (±5 kb) with DEK siRNA-mediated knockdown and control **f,** MNase accessibility assay following CX-4945 treatment in control and DEK knockdown SK-Mel-103 cells; DNA recovered from supernatant and pellet fractions was resolved by agarose gel electrophoresis.

To directly assess the impact of CK2-dependent DEK regulation on chromatin organization, we monitored nuclear morphology and nuclease sensitivity. Treatment with the CK2 inhibitor CX-4945 induced methuosis-like cytoplasmic vacuolization and nuclear condensation, as indicated by a ∼42% reduction in DAPI-positive nuclear area (p < 0.001) together with decreased global MNase accessibility in SK-Mel-103 cells (**Fig. 7a and c, Extended data Fig. 8b**). Strikingly, within the same cellular context, siRNA-mediated DEK depletion substantially mitigated CX-4945-induced chromatin condensation, reducing nuclear compaction by approximately 60% and completely restoring MNase sensitivity, while having minimal impact under basal conditions (**Fig. 7c and f**).

ATAC-seq further highlighted the phosphorylation-dependent nature of this process. Under basal conditions (DMSO), DEK depletion caused only modest changes in global MNase sensitivity and local ATAC-seq signals (chromatin accessibility) at DMSO-specific DEK occupied regions. By contrast, upon CK2 inhibition, DEK knockdown led to dramatic reduction of local ATAC-seq accessibility at CX-4945-unique DEK binding sites, accompanied by a marked increase in MNase sensitivity. Comparable patterns were observed in HeLa S3 cells, where S287/S288 remained unphosphorylated, further linking DEK hypophosphorylation to the stabilization of compact chromatin states (**Fig. 7e and f**).

Together, these findings reveal that hypophosphorylated DEK reinforces transcriptional repression through chromatin compaction and maintenance of local accessibility. CK2-dependent phosphorylation thus operates as a molecular switch that modulates DEK’s chromatin-binding capacity, emphasizing that its activity cannot be inferred from expression levels alone. This positions DEK as a context-dependent, phosphorylation-responsive regulator of chromatin architecture, providing a conceptual framework for the development of context-specific therapeutic strategies targeting DEK regulation in diverse tumor types, with particular relevance to melanoma progression.

## Discussion

Our work identifies the DEK oncoprotein as a phosphorylation-sensitive regulator of chromatin organization whose genomic engagement is strongly cell-type specific. By integrating ChIP–seq, ATAC–seq, phosphoproteomics and transcriptional profiling across primary, transformed and melanoma cells, we show that DEK protein abundance alone is a poor predictor of its chromatin association and transcriptional impact. Instead, CK2-dependent phosphorylation, particularly at serines 287 and 288 within the C-terminal DNA-binding region, dictates whether DEK is immobilized on chromatin to stabilize compact repressive domains or exists in a more mobile, weakly chromatin-tethered state, potentially associated with transcriptionally active programs. This framework reconciles the widely observed upregulation of DEK in cancer with the striking variability in its chromatin occupancy and regulatory consequences across different cellular contexts.

The distribution of DEK across chromatin compartments has long been debated. Hu *et al*. reported DEK enrichment in euchromatin without detectable co-localization with the heterochromatin marker HP1α, whereas Kappes *et al*. demonstrated direct DEK–HP1α interaction and implicated DEK in heterochromatin maintenance, and genome-wide analyses in K562 and U937 cells found preferential DEK binding at chromatin marked by H3K4me3, H3K27ac and H3K9ac, consistent with an association with open chromatin ^17, 38, 47^. Our ChIP-seq data in HeLa S3 and melanoma cells align with these latter observations, with DEK predominantly localized to promoters and enhancers marked by H3K4me3 and H3K27ac, yet our functional analyses reveal that this broadly euchromatic localization can support both transcriptional repression and activation in a location-dependent manner. To dissect how this distribution translates into distinct regulatory outcomes, we examined DEK function at the genome-wide level. DEK shows a pronounced preference for promoter-proximal chromatin, while also engaging in a discrete set of distal regulatory elements. Within this promoter-centric binding landscape, integration of occupancy, accessibility and expression data indicates that DEK most often constrains transcription at its promoter-bound targets, with approximately two-thirds of genes that change upon DEK loss being upregulated. Metaprofiles of chromatin accessibility further reveal that this repression is not achieved through indiscriminate chromatin closure. At promoters, DEK sharpens local architecture by maintaining a focused accessible region at its binding site while limiting excessive opening of surrounding sequences. By contrast, at distal intergenic elements DEK contributes to maintaining an open chromatin configuration, and its loss is accompanied by a more uniform reduction in accessibility that, for a subset of linked genes, parallels decreased expression. Thus, DEK’s impact on chromatin and transcription is intrinsically context dependent and may be executed by differentially modified DEK subpopulations. This duality provides a mechanistic explanation for why DEK has been described both as a heterochromatin-associated repressor and as a facilitator of transcription in euchromatic regions ^12, 28, 43, 48–50^.

At the DNA level, early reports of apparent sequence-specific binding at HIV-2 pets and MHC promoters have been revised by biochemical work showing that DEK primarily recognizes DNA in a structural manner, favoring supercoiled and four-way junction DNA, akin to HMG-box proteins that bend DNA and modulate chromatin architecture ^24, 51, 52^. Despite this predominantly structural binding mode, DEK exhibits gene-specific associations. ChIP–seq in HeLa S3 and FM232 cells, together with previous studies in other systems ^38^, identify TOP1 (topoisomerase I; **Extended data Fig. 9c**) as a prominent and conserved DEK target. TOP1 resolves DNA torsional stress, and DEK binding at its promoter suggests functional interplay, consistent with observations in *Arabidopsis*, where DEK and TOP1 co-regulate flowering ^53, 54^. In addition, DEK co-localizes with Pol II and GC-rich motifs in our datasets, implicating it in transcriptional activation at a subset of promoters. However, whether DEK engages these loci through direct DNA recognition or predominantly via recruitment by transcription factors such as ZNF770 or KLF5 remains unresolved. Taken together, these prior studies and our genome-wide analyses support a parsimonious interpretation in which DEK functions primarily as a chromatin architectural regulator whose apparent euchromatin preference and diverse transcriptional outcomes are dictated by genomic location, interacting partners, local chromatin state and modification pattern.

Having established this locus-level model, we next asked to what extent DEK’s architectural functions are conserved across cell types or are themselves subject to cell-type-specific regulation. We further demonstrate that this dual functionality is embedded in a framework of strong cell-type specificity. Although DEK shows a broadly conserved preference for promoter-proximal occupancy in primary melanocytes and HeLa S3 cells, the networks of DEK-bound genes diverge substantially between normal and transformed contexts. In primary melanocytes, DEK preferentially occupies genes involved in transcriptional and chromatin regulation, whereas in HeLa cells it is enriched at genes related to signaling and transport. This divergence suggests that lineage-defining transcription factors, local chromatin accessibility and cofactor composition jointly determine which genomic regions are competent for DEK engagement.

The particularly striking case of melanoma cell lines illustrates how these determinants intersect with post-translational regulation. In SK-Mel-103, DEK is highly expressed and biochemically tethered to chromatin, yet largely “ChIP-invisible” under basal conditions, underscoring that conventional peak-based ChIP analyses may underestimate DEK occupancy when interactions are highly dynamic, organized at three-dimensional loop anchors or embedded within phase-separated chromatin condensates ^55–57^. In these melanoma cells, DEK is extensively phosphorylated at CK2 sites, a modification known to weaken DNA binding, raising the possibility that hyperphosphorylated DEK partitions preferentially into highly dynamic condensates with limited stable DNA anchoring, supported by sucrose density gradients, native gels and the lack of chromatin campaction upon hyperphosphorylation in SV-40 minichromosomes. Consistent with this idea, even mild pharmacological perturbation, including DMSO exposure and CK2 inhibition, can shift DEK from a weakly DNA-anchored, condensate-like state into more stably chromatin-bound configurations that are more readily detected by peak-based ChIP analysis. More broadly, phase separation of chromatin regulators such as HP1 has emerged as a general mechanism for assembling repressive nuclear compartments whose material properties are tuned by phosphorylation and nucleic acid binding, suggesting that DEK may act as a phosphorylation-sensitive client or scaffold within such condensates ^57–59^. These observations cleary warrant further investigations, yet prompted us here to investigate in greater detail how CK2-dependent phosphorylation tunes DEK’s chromatin-binding capacity and thereby its functional partitioning between chromatin compartments.

A central mechanistic insight from this study is that CK2-mediated phosphorylation of DEK’s C-terminal DNA-binding region tunes its chromatin-binding capacity and thereby its functional partitioning across the genome. In line with biochemical work showing that CK2 phosphorylation in this region decreases DEK-DNA and DEK-chromatin affinity while promoting multimerization ^42^, our phosphoproteomic and Phos-tag analyses identify extensive phosphorylation of DEK at CK2 consensus sites, including S287/S288, in SK-Mel-103 cells, and CK2 inhibition reduces this phosphorylation and redistributes DEK from broad enhancer-like euchromatic regions towards H3K9me3-enriched repressive chromatin, with increased promoter-proximal occupancy and concordant transcriptional downregulation at a subset of targets. Chromatin compaction can in principle be achieved by nucleosome bridging and crosslinking, loop extrusion, histone modification and ATP-dependent remodelling ^60–64^; although DEK lacks known enzymatic activity ^3^, cryo-EM structures and functional studies show that DEK dimers engage nucleosomes via an N-terminal histone-binding/pseudo-SAP module and a C-terminal basic tail, introduce ∼30° kinks into linker DNA, interact with HP1α and cooperate with H3K9me3-marked heterochromatin, and can enhance PRC2-mediated H3K27 trimethylation, while heterologous expression of DEK in bacteria, which lack canonical nucleosomes, is nevertheless sufficient to compact DNA ^27, 28, 45^. By mapping S287/S288 as CK2-regulated sites and linking their phosphorylation state to altered chromatin association and MNase sensitivity, we provide in-cell evidence that C-terminal phosphorylation acts as a tunable “off-switch” for these architectural activities: under hypophosphorylation conditions, DEK is preferentially redistributed towards H3K9me3-enriched regions where it has been reported to interact with HP1α, and DEK depletion then causes a stronger increase in MNase sensitivity than depletion of predominantly hyperphosphorylated DEK, consistent with a more prominent contribution of hypophosphorylated DEK to chromatin compaction. Thus, a single kinase-substrate pair can toggle DEK between chromatin-immobilized, compaction-prone and more mobile, euchromatin-associated modes without changing DEK expression levels.

More broadly, the CK2-DEK axis illustrates how phosphorylation can reprogram non-histone chromatin proteins, flipping DEK between chromatin-compacting and more permissive modes and likely mirroring analogous post-translational “mode switches” in other architectural factors. This switch is further constrained by oncogenic context, as only NRAS^Q61R^ melanoma cells (SK-Mel-103), but not BRAF^V600E^ lines (UACC-87) with comparable DEK expression, show robust DEK redistribution and gene repression upon CK2 inhibition, suggesting that phospho-DEK patterns and DEK chromatin-binding signatures could help identify tumor settings in which DEK-mediated chromatin reprogramming is therapeutically exploitable. CK2 inhibition also reveals broader functions of DEK beyond chromatin, increasing salt-resistant chromatin association while redistributing DEK to the cytosol and extracellular space and enhancing its RNA-binding capacity. Consistent with this, our transcriptomic analysis shows that DEK depletion leads to strong upregulation of multiple SNORD116 small nucleolar RNAs among the top induced genes (**Fig 2a**), and together with previous work implicating SNORD116 in alternative splicing via FOX2 and DEK itself in genome-wide splicing control and ribosome biogenesis, this suggests that phosphorylation-sensitive DEK coordinates chromatin regulation with RNA processing pathways ^65–69^. Together, these features support a view of DEK as an integrative regulator at the intersection of kinase signalling, chromatin architecture, RNA metabolism and extracellular communication.

From a therapeutic perspective, targeting DEK phosphorylation, for example by using CK2 inhibitors, may offer a means to reprogram chromatin and RNA-processing networks in melanoma, particularly in tumors that retain a hypophosphorylation-competent DEK pool. A key challenge will be to define robust phospho-DEK and chromatin signatures that predict such dependency and to determine how selectively modulating the CK2-DEK axis can be achieved without perturbing other essential kinase targets and RNA-processing pathways.

## Materials and Methods

Supplementary Table 1 list all cell lines, Supplementary Table 2 lists antibodies used, Supplementary Table 3 list all ChiP-seq experiments performed, Supplementary Table 4 list enzymes used, Supplementary Table 5 lists oligonucleotides, Supplementary Table 6 list chemicals and Supplementary Table 7 list all kits used in this study.

### Cell lines, primary cells and cell viability assays

HeLa S3 cells, melanoma cell lines SK-Mel-19, SK-Mel-29, SK-Mel-103, UACC-62, UACC-257, and full-length DEK-deficient derivative 1E7 line were maintained in DMEM supplemented with 10% fetal bovine serum (FBS; Gibco), Penicillin-Streptomycin 100 X solution (Hyclone) at 37 °C in a humidified incubator with 5% CO_2_. Primary foreskin melanocytes (FM232) were cultured 254CF medium (Gibco) supplemented with human melanocyte growth supplement (HMGS-2, PMA free, Gibco), Penicillin-Streptomycin 100 X solution (Hyclone). Other primary melanocytes and primary mammary epithelial cells commercially available from Procell were cultured in complete medium specific to cells as supplied by Procell.

For cell viability flow cytometry, cells were treated with CX-4945 for 24 hours at the concentration of 5, 10, 20, and 40 µM, respectively. Non-treated cells served as negative control and DMSO-treated cells served as vehicle control. Cell culture medium was aspirated and collected into a 15 mL conical tube. After two wash steps with 1X PBS and trypsinization, cells were resuspended in the supernatants collected previously followed by one more wash with 1X PBS. To fix the cells, they were centrifuged at 200 x g for 3 minutes and then incubated with ice-cold 70% ethanol for 2 hours on ice. Cells were subsequently centrifuged at 200 x g for 3 minutes and washed with 1 mL ice-cold PBS again. For staining, cells were subjected to 500 µL propidium iodide (PI) treatment at 37 °C for 30 minutes in the dark. Afterwards, cells were centrifuged at 200 x g for 3 minutes, resuspended with 1X PBS, and filtered into round bottom Fluorescence-Activated Cell Sorting (FACS) tubes on ice prior to flow cytometry analysis. The excitation wavelength for generating Forward Scattering (cell size) (FSC) and Side Scattering (cell granularity) (SSC) was 488 nm.

To detect apoptosis, ABflo® 488 Annexin V/PI Apoptosis Detection Kit (Abclonal) was used according to the manufacturer’s instructions. Briefly, cells were harvested, washed twice with cold PBS, and resuspended in 1×binding buffer at a concentration of 1×10^6^ cells/mL. A 100 µL aliquot of the cell suspension was transferred to a flow cytometry tube, and 5 µL of Annexin V-FITC and 5 µL of propidium iodide (PI) were added. The mixture was gently vortexed and incubated in the dark at room temperature for 15 minutes. After incubation, 400 µL of 1 × binding buffer was added to each tube, and the samples were analyzed immediately using a flow cytometer. Single-stained cells (PI-only or Annexin V-only) were used to adjust compensation. A DNA ladder assay was used to detect cell apoptosis by examining the fragmented genomic DNA on a agarose gel. Cells were lysed in 0.5 mL detergent buffer (10 mM Tris, pH 7.4, 10 mM EDTA, 0.2% Triton X-100), vortexed, and incubated on ice for 30 min. The lysate was centrifuged at 27,000 × g for 30 minutes, and the supernatant was collected. DNA was precipitated by adding 100 µL of ice-cold 5 M NaCl and was vortexed, followed by ethanol precipitation. The precipitated DNA pellets were pooled by resuspending in 400 µL of extraction buffer (10 mM Tris, 5 mM EDTA). To remove RNA, 1 µL of RNase I (Thermo Scientific) was added, and the samples were incubated at 37°C for 5 hours. Protein digestion was carried out by adding 1 µL of proteinase K (800 units/mL, NewEngland Biolabs), followed by overnight incubation at 65°C. DNA was extracted again using DNA extraction solution (Solarbio) according to manufacturer’s instruction and precipitated with ethanol. The resulting DNA was resuspended in nuclease-free H2O and separated on an agarose gel for visualization of apoptotic DNA ladder pattern.

### DEK depletion, knockdown, and lentiviral transduction

The full-length DEK-deficient HeLa S3 clone 1E7 was generated previously by TALEN (transcription activator-like effector nucleases)-mediated genomic disruption ^45^. For transient knockdown, cells were transfected with *DEK*-targeting or non-targeting control siRNAs using Lipofectamine 3000 (Thermo Fisher Scientific) according to the manufacturer’s instructions. Briefly, siRNA and 5 µl Lipofectamine 3000 were each diluted in 125 µl Opti-MEM, incubated separately for 5 min at room temperature, combined and incubated for a further 20 min to allow complex formation, and then added directly to cells. When cells reached approximately 80% confluency, 40 µM CX-4945 (Silmitasertib; MCE) dissolved in DMSO, or an equivalent volume of DMSO as vehicle control, was added to parallel cultures. After 24 h incubation at 37°C in a humidified atmosphere with 5% CO_2_, cells were collected for chromatin fractionation, MNase digestion, ChIP–seq or other downstream analyses as indicated. Lentiviral transduction of SK-Mel-103 cells with pTRIPZ plasmids expressing GFP, GFP-full length DEK, or a GFP-full-length DEK mutant with serines 227, 230, 231, 232, 243, 244, 251, 301, 303, 306, 307 mutated alanine (S-11-A) was carried out as described in^19^. Briefly, a second generation lentiviral system that allows for inducible gene expression was employed following the procedures of the RNAi consortium (https://www.addgene.org/protocols/plko/#E). On the first day, approximately 3× 10^5^ HEK-293FT packaging cells were plated on 6 cm tissue culture dishes with 6 mL of low-antibiotic growth medium (DMEM + 10 % FBS + 0.1 x Pen/Strep) and incubated for 24 hours. The plasmid mixtures were prepared as follows: packaging plasmid (psPAX2, 900 ng), envelope plasmid (VSV-G, 100 ng), pTRIPZ vector (1 μg), P3000™ reagent (10 μL) and Opti-MEM medium to total volume of 30 μL. Six μL of Lipofectamine 3000 transfection reagent were diluted with Opti-MEM medium to a final volume of 90 µL. The plasmid mixture was then added dropwise to the diluted Lipofectamine 3000 transfection reagent and incubated at RT for 25 minutes before being transferred to the packaging cells. The medium was replaced with 6 mL high FBS medium (DMEM + 30 % FBS + 1 x Pen/Strep) 18 hours post transfection. Viral particles were harvested roughly 40 hours after transfection, filtered through 0.45 µm PVDF filters into a 15 ml tube, and mixed with 6 mL of normal medium containing 8 µg/mL polybrene. The packaging cells were replenished with 6 mL of high FBS medium for a secondary viral harvest 24 hours later. The target cells, which had been seeded one day prior in 10 cm tissue culture plates, were incubated with the virus-containing solution for 6 hours before replacement with normal medium. After two rounds of transduction, the cells were selected with 1 µg/mL puromycin (InvivoGen) for at least 48 hours. Established pTRIPZ cell lines with constitutive puromycin-resistance marker were maintained in medium supplemented with 1 µg/mL Puromycin (InvivoGen). For induction of GFP-DEK expression, cells were cultured with medium containing 1 µg/mL Doxycycline (Beyotime) for 24/48 hours.

### Cell, chromatin fractionation and immunoprecipitation

To resolve DEK across different cellular compartments, we separated cytosolic, soluble nuclear and chromatin-bound fractions by stepwise subcellular fractionation. Cells were washed twice with ice-cold PBS, scraped with a rubber scraper and pelleted by centrifugation at 500 x g for 5 min. Pellets were rapidly rinsed once in ice-cold hypotonic buffer (20 mM HEPES pH 7.6, 20 mM NaCl, 5 mM MgCl_2_) and collected again at 500 x g for 5 min, then resuspended in 5 ml hypotonic buffer and transferred to a Dounce homogenizer. After a 10-min incubation on ice, cells were disrupted by 10 strokes with a loose pestle and the homogenate was centrifuged at 500 x g for 5 min to yield the cytosolic supernatant. The pellet was resuspended in hypotonic buffer containing 0.5% NP-40 to extract soluble nuclear proteins, followed by centrifugation at 1,000 x g for 5 min. Chromatin-associated proteins were then sequentially extracted by resuspending the remaining pellet in extraction buffer (20 mM HEPES pH 7.6, 300 mM sucrose, 0.5 mM MgCl_2_) supplemented with increasing NaCl concentrations (100 mM, 250 mM and 450 mM; 15 min each), and collecting the corresponding salt-soluble chromatin fractions after centrifugation at 1,000 x g for 5 min. The final pellet was solubilized in 2% SDS to obtain the residual chromatin fraction. All buffers were freshly supplemented with 1 mM Na_3_VO_4_, 10 mM NaF and 1× cOmplete protease inhibitor cocktail (Roche). The distribution of DEK and marker proteins in each fraction was assessed by immunoblotting.

For immunoprecipitations, the nuclear extracts were incubated with either 1 µg antibody targeting a protein of interest or 1 µg of unspecific IgGs in a tube rotator at 4 °C overnight. The second day, 20 µL of pre-equilibrated magnetic protein A+G beads (Beyotime) slurry was added to the antibody-antigen complex and incubated for another 2 hours at 4 °C. The beads were washed once with 1 mL buffer B (20 mM HEPES, pH 7.6, 450 mM NaCl, 0.5 % IGEPAL, 1 mM Na_2_VO_3_, 10 mM NaF, 1 × cOmplete protease inhibitor cocktail, 1 mM PMSF) and twice with 1 mL buffer B without IGEPAL. For immunoprecipations of GFP-tagged proteins, nuclear extracts were incubated with 20 µL of equilibrated GFP-Trap (Chromotek) beads or agarose beads as control. The beads were then resuspended in 20 µL of 2X Laemmli loading buffer and boiled at 95 °C for 5 minutes for further SDS-PAGE analysis.

### Micrococcal nuclease (MNase) digestion

MNase digestion was used to assess global chromatin accessibility using either a time course setting or a differential extraction method modified from^40^. For time course experiments, adherent cells were washed twice with ice-cold PBS, scraped and pelleted at 500 x g for 5 min. Approximately 1.2 × 10^7^ cells per condition were washed once in 10 ml hypotonic buffer (20 mM HEPES pH 7.6, 20 mM NaCl, 5 mM MgCl_₂_) freshly supplemented with 1 mM Na_3_VO_4_, 10 mM NaF, 1× cOmplete protease inhibitor cocktail (Roche) and 1× PhosSTOP (Beyotime), collected again at 500 x g and resuspended in 5 ml hypotonic buffer. After gentle inversion, suspensions were Dounce-homogenized (10 strokes with a loose pestle), incubated on ice for 10 min and IGEPAL (Sigma) was added dropwise to 0.5% with gentle mixing. Following a further 5-min incubation, nuclei were pelleted at 500 x g for 5 min and washed twice in extraction buffer (20 mM HEPES pH 7.6, 100 mM NaCl). Nuclei were resuspended in extraction buffer supplemented with 2 mM CaCl_2_ to 2 × 10^6^ cells per 100 µl, and 100-µl aliquots were digested at 25 °C with micrococcal nuclease (2,000,000 U ml⁻¹; New England Biolabs) for 0, 2, 4, 8, 16 or 32 min. Reactions were stopped by addition of EDTA to 8 mM, and digested nuclei were spun at 16,000 x g for 2 min. Supernatants were adjusted to 1% SDS; highly viscous samples (t = 0) were briefly sonicated. Samples were treated with proteinase K (800 U ml⁻¹; New England Biolabs; 1 µl per reaction) at 55 °C for 30 min and DNA was extracted using DNA extraction reagent (Solarbio) followed by ethanol precipitation. Purified DNA was resolved on agarose gels to visualize mono- and oligonucleosome ladders. Differential extraction was carried out as described before ^23^ by washing cells with ice-cold PBS, scraping, and lysis in HNB buffer (500 mM sucrose, 15 mM Tris pH 7.5, 60 mM KCl, 0.25 mM EDTA, 0.5 µM spermidine, 0.15 µM spermine, 1 mM DTT, 10 mM NaF, 1 mM NaV, 1x Complete.) containing 0.5% NP-40. Resulting nuclei were pelleted and washed in Nuclear Buffer (20 mM Tris pH 7.5, 70 mM NaCl, 20 mM KCl, 5 mM MgCl₂, 1 mM PMSF, 1 mM DTT, 10 mM NaF, 1 mM NaV, 1x Complete) and DNA concentration was determined by A_260_. Chromatin (500 µg/mL) was digested with 75 U MNase/50 µg chromatin in the presence of 2 mM CaCl₂ at 22°C. Aliquots were taken at 0, 1, 2, and 4 min, and reactions were stopped with 2 mM EDTA. Samples were centrifuged (5 min, 16,000 x g, 4°C). The supernatant (S1, easily accessible chromatin) was collected. The pellet was resuspended in 8 mM EDTA, incubated on ice, and re-centrifuged. The resulting supernatant (S2, inactive, dense chromatin) and final pellet (P, matrix-associated chromatin) were collected. For immunoprecipitations of DEK from chromatin fractions obtained from the SK-MEL 103 cell line, a chromatin fractionation was carried out (volume per fraction ∼600 μL), S2 and P fractions were adjusted to nuclear buffer conditions, and NaCl concentration in all fractions was adjusted to 550 mM. The samples were mixed and incubated on ice for 10 minutes. After a centrifugation step (10 min, ∼15700 x g, 4 °C), the supernatants were collected, divided (1x IP, 1x control) and the volume was adjusted to 1 mL with nuclear buffer. For the precipitation, 1 μg of the polyclonal anti-DEK antibody K877 or 1 μg rabbit IgGs were added and the samples were incubated in an overhead shaker (1 h, 4 °C), followed by addition of 15 μL of pre-equilibrated Protein A Sepharose and three wash steps with 1 mL NTEN buffer (20 mM Tris pH 7.5, 150 mM NaCl, +/- 1 mM EDTA (for dephosphorylation: NTEN 150 without EDTA), 0.5% NP-40, 1x Complete). If a GFP-trap (Chromotek) was used instead of IgGs and Protein A-coupled Sepharose, the addition of antibodies was omitted. Instead of Protein A Sepharose, 10 μL of GFP-trap (control: 10 μL bab Sepharose) was added and the procedure continued as described above. For dephosphorylation with λ-phosphatase, the samples were centrifuged, the supernatant was removed, and 50-100 μL of phosphatase buffer and 1200 U of λ-phosphatase were added. The samples were dephosphorylated in a thermal shaker (1 h at 30°C / 40 min at 37°C) and subsequently washed three times with NTEN 150 (Protein A Sepharose: 5 min at 1000 x g, GFP-trap: 5 min at 2700 x g, 4°C). To assess DEK-HP1 interactions, 3 μL of interaction mix (10 μg BSA (10 mg/mL), 30 μL NTEN 150, 20 μL HP1 overexpression in *E. coli*) was added to the samples (1 mL NTEN 150 with Protein A Sepharose beads/GFP-trap and bound DEK). The samples were incubated on an overhead shaker (1 h, 4 °C) and washed three times with NTEN 150. The pellet was resuspended in 1x Laemmli sample buffer and boiled for five minutes at 95 °C.

### Phos-tag SDS-PAGE and immunoblotting

Phosphorylated DEK species were resolved using Phos-tag SDS–PAGE. For Phos-tag gels, 50 µM Phosbind acrylamide (APExBIO) and 100 µM MnCl_2_ were added to the separating gel before polymerization. An EDTA-free prestained protein marker (APExBIO) was used as a standard. After electrophoresis, gels were incubated twice in transfer buffer containing 5 mM EDTA (10 min each) and once in EDTA-free transfer buffer (10 min) before blotting. For immunoblotting, proteins were transferred to PVDF membranes (Millipore Sigma). Membranes were activated by brief soaking in methanol, rinsed in distilled water and equilibrated in 1× transfer buffer. Transfer was performed either by semi-dry blotting (Biometra; 12 V, 1 h) or wet transfer (Bio-Rad; 140 mA for 1 h or 20 mA overnight). Membranes were blocked in 5% skimmed milk in PBS-0.1% Tween-20 (PBS-T) for standard proteins, or 5% BSA in PBS for detection of phosphorylated proteins. Primary antibodies were diluted in primary antibody dilution buffer (Beyotime) and incubated with membranes for 1 h at room temperature or overnight at 4 °C with agitation. After three washes in PBS-T (10 min each), membranes were incubated with HRP-conjugated secondary antibodies (Beyotime; 1:1,000 in primary antibody dilution buffer) for 1 h at room temperature, washed four times in PBS-T (5 min each) and developed using the BeyoECL Moon kit (Beyotime) according to the manufacturer’s instructions. Chemiluminescence was recorded on a ChemiDoc imaging system (Bio-Rad).

### Mass spectrometry and phosphosite identification

Immunoprecipitated proteins were separated by SDS–PAGE and visualized by Coomassie blue staining. Bands at the expected molecular weight were excised and diced into ∼1 mm^3^ cubes, transferred to 1.5-ml tubes and washed in HPLC-grade water (10 min, room temperature (RT) followed by 50 mM ammonium bicarbonate (AB; 10 min, RT). Gels were de-stained twice with 50% acetonitrile (ACN) in 50 mM AB at 37 °C for 20 min and dehydrated in 100% ACN for 5 min. For reduction and alkylation, gel pieces were incubated with 10 mM dithiothreitol (DTT) in 100 mM AB at 56 °C for 40 min and then with 55 mM iodoacetamide (IAA) in 100 mM AB at RT for 45 min in the dark. After removal of excess reagent and brief dehydration, proteins were digested in-gel overnight at 37 °C with sequencing grade modified trypsin (Promega; 1 µg in 100 µl 50 mM AB). The following day, peptide-containing supernatants were collected by brief centrifugation and peptides were further extracted from the gel pieces by sequential incubation in HPLC-grade water (37 °C, 20 min), 5% formic acid (FA) in 50% ACN (two rounds, 100 µl, 37 °C, 20 min each) and finally 100% ACN (5 min, RT). All extracts were pooled and dried to completeness in a SpeedVac vacuum concentrator (Thermo Scientific). Dried peptides were resuspended in 2% ACN/0.1% FA and desalted on Monospin C18 columns (GL Sciences) according to the manufacturer’s instructions. Briefly, columns were activated with 100 µl 100% ACN and equilibrated with 0.1% trifluoroacetic acid (TFA), with solutions passed through by centrifugation at 5,000g for 1 min. Peptide solutions were loaded, washed with 0.1% TFA and eluted with 0.1% TFA/60% ACN. Eluates were dried in a SpeedVac and reconstituted in 5% ACN/0.1% FA prior to LC–MS analysis.Raw Orbitrap data files were inspected in Xcalibur (Thermo Fisher Scientific) and searched using Mascot Daemon against the appropriate UniProt database. Searches were performed with trypsin (specific, allowing for the modified trypsin cleavage rules) as the protease, carbamidomethylation of cysteine (C) as a fixed modification and phosphorylation, N-terminal acetylation and methionine (M) oxidation as variable modifications. Peptide-spectrum matches and phosphosite assignments were manually validated.

### Microarray gene expression profiling

Transcriptome profiling of HeLa S3 wild-type and 1E7 cells was performed using the GeneChip Human Transcriptome Array 2.0 (HTA 2.0; Affymetrix). Total RNA was isolated using the RNeasy Mini Kit (QIAGEN) according to the manufacturer’s instructions and quantified and quality-controlled before hybridization. cDNA synthesis, labelling and hybridization to HTA 2.0 arrays were carried out at the IZKF Chip Facility, RWTH Aachen University (Aachen, Germany), following the Affymetrix protocols. Raw CEL files were processed in Transcriptome Analysis Console (Thermo Fisher Scientific) using robust multi-array averaging for background correction and normalization, and differentially expressed genes were identified with the software’s built-in statistical workflow.

### ATAC-seq

Genome-wide chromatin accessibility was profiled by Assay for Transposase-Accessible Chromatin with sequencing (ATAC-seq). ATAC-seq was performed by Active Motif (Shanghai, China) on HeLa S3 wild-type and 1E7 cells, as well as SK-Mel-103 cells with or without CX 4945 treatment. Cells were trypsinized, resuspended in freezing medium (FBS:growth medium:DMSO = 5:4:1), frozen at a controlled cooling rate and shipped on dry ice. Library preparation and sequencing were carried out by Active Motif using their standard Tn5-based ATAC-seq protocol and Illumina sequencing.

Sequencing reads were trimmed and aligned to the human reference genome (GRCh38/hg38); low-quality, duplicate and mitochondrial reads were removed, and peaks of accessible chromatin were called using standard ATAC-seq pipelines. For metaprofiles, normalized ATAC–seq coverage (reads per million) from bigWig files was extracted in fixed-size bins around transcription start sites or DEK ChIP-seq peaks using deepTools computeMatrix, and the mean signal per bin across all regions was plotted using plotProfile.

### ChIP-seq

Cells were crosslinked in 1% formaldehyde for 10 min at room temperature, quenched with 125 mM glycine, washed in cold PBS and lysed in a high-salt lysis buffer. Nuclei were sonicated in a Bioruptor to yield chromatin fragments of ∼200-500 bp, as confirmed by agarose gel electrophoresis. After clarification, chromatin was pre-cleared with protein A/G magnetic beads and incubated with antibodies (anti-DEK, anti-H3K9me3, anti-H3K27ac, Abcam; IgG control, Beyotime) at 4 °C overnight. Immune complexes were captured with magnetic beads, washed through a series of low-salt, high-salt and LiCl buffers, and eluted in SDS buffer. Crosslinks were reversed by overnight incubation at 65 °C, followed by RNase A and proteinase K digestion. DNA was purified using spin columns and quantified. ChIP-seq libraries were prepared and sequenced using single-end 50 bp sequencing (SE50) on the BGISEQ-500 platform (BGI, Wuhan, China).

Sequencing quality was assessed with FastQC, and FASTQ files were converted and processed using the FASTQ groomer where necessary. Clean reads were aligned to the human reference genome GRCh38/hg38 using Bowtie2, and unmapped reads, PCR duplicates, low-quality reads were removed. Peaks were called with MACS2 (q < 0.01) using matched input controls. Three independent ChIP-seq experiments per condition were analysed, and irreproducible discovery rate (IDR) analysis was used to define high-confidence peaks. Peaks were annotated to promoters, gene bodies and distal intergenic regions using ChIPseeker. Motif enrichment analyses within DEK peaks were performed with MEME-ChIP. Differentially bound sites were addressed by R package Diffbind (v2.10.0) (Rory Stark, 2022). Kyoto encyclopedia of genes and genomes (KEGG) pathway enrichment analysis and gene ontology enrichment analysis were performed by the online platform ShinyGO (v0.75, http://bioinformatics.sdstate.edu/go/).

Selected ChIP-seq peaks were validated by quantitative PCR. ChIP and corresponding input DNA were analysed by SYBR Green-based qPCR on a QuantStudio 5 Real-Time PCR System (Applied Biosystems) using primers spanning ChIP-seq peak summits and control regions as indicated in the figure legends. Relative enrichment was calculated using the percent-input method, in which input Ct values were adjusted for dilution and ChIP signals expressed as a percentage of input; where indicated, fold enrichment over IgG control was also determined.

### RNA extraction and quantitative PCR (qPCR)

To isolate RNA, TRIzol reagent (Beyotime) was added directly onto the cell culture dish. The lysate was pipetted up and down several times to thoroughly lyse the cells and then kept at room temperature for 5 minutes. Two hundred microliters of chloroform were added to 1 mL TRIzol, followed by vigorous vortexing for 15 seconds. The lysate was kept at room temperature for another 5 minutes. After centrifugation at 12,000 x g for 15 minutes, the upper aqueous phase containing RNA was transferred to a new EP tube, followed by the addition of 0.5 mL isopropanol per 1 mL TRIzol and 2 μL glycogen. The RNA was precipitated at -20 °C overnight and then centrifuged at 13,000 x g for 15 minutes at 4 °C. The pellet was washed once with 70% ice-cold ethanol, which was followed by centrifugation at 7,500 x g for 5 minutes. RNA pellets were air dried at room temperature and dissolved in 30 μL of DEPC water, followed by an RNA Cleanup procedure using RNeasy Kit (QIAGEN) following the manufacturer’s instructions and eluted with 50 μL of nuclease free water.. The concentration and purity of the obtained RNA were determined by NanoDrop (Thermo Scientific). Reverse transcription polymerase chain reaction (RT-PCR) was performed by using one μg RNA mixed with Oligo(dT)15 (500 μg/mL) primers in a total volume of 5 μL, followed by heating at 70 °C for 5 minutes. Then GoScript™ 5X Reaction Buffer, MgCl2, PCR Nucleotide Mix, RNasin® Ribonuclease Inhibitor, and GoScript™ Reverse Transcriptase were added and subjected to amplification. The resulting cDNA products were diluted 50-fold with ddH_2_O for subsequent qPCR analysis. The reactions were conducted on the Applied Biosystems QuantStudio™ 5 Real-Time PCR system with using standard setting reccomened for the GoTaq® qPCR Master Mix. Relative gene expression was assessed using the comparative delta-delta Ct method (2–ΔΔCt)

For ChIPed DNA, enrichment was measured either by the percentage of input method or the fold enrichment method. The percentage of input method firstly adjusts the input to 100%, followed by comparison of DNA pulled down relative to the amount of input, using the formula below: percentage of Input = 100*2^ (Adjusted input - Ct (IP)*Adjust Input = Ct Input - log 2(Dilution factor). Alternatively, fold-enrichment was calculated by comparing the amount of DNA pulled down in the ChIP sample to that in a negative control (IgG), using the formula: Fold-Enrichment = 2^(Ct_IgG – Ct_ChIP). The amplicon size and primer specificity were verified through melt curve analysis in combination with gel electrophoresis of the qPCR amplification products.

### Immunofluorescence microscopy

For immunofluorescence, cells grown on glass coverslips were fixed in 4% paraformaldehyde for 10 min, permeabilized with 0.5% Triton X-100 and blocked in 1% BSA/22.52 mg/mL glycine/PBS-T for 30 minutes. Coverslips were incubated with primary antibodies diluted in 1% BSA/PBS-T for 1 h at room temperature or overnight at 4 °C, washed and incubated with Alexa Fluor-conjugated secondary antibodies for 1 h. Nuclei were counterstained with DAPI (Beyotime) and coverslips were mounted in anti-fade medium. Images were acquired on a confocal microscope (Zeiss) with identical settings within each experiment and analyzed in Fiji/ImageJ.

### Statistical analysis

Unless otherwise indicated, all experiments were performed with at least three independent biological replicates. Data are presented as mean ± s.e.m. or mean ± s.d. as specified in the figure legends. Statistical analyses were performed in GraphPad Prism. Comparisons between two groups were made using two-tailed unpaired Student’s t-tests, and multiple comparisons were analyzed by one-way ANOVA. P values < 0.05 were considered statistically significant.

## Acknowledgments

We are grateful to Bernd Denecke (IZKF, RWTH Aachen University) for conducting microarray assays, Philippe Collas for the pH3.3-EGFP construct, Chutian Chen for conducting BGIS, Anja Deutzmann and Elisa Ferrand-May for conducting experiments leading to Fig. 4d.

## Funding

DFG (Deutsche Forschungsgemeinschaft) KA 2799/1 (FK)

RDF (Research Development Fund), Xi’an Jiaotong-Liverpool University, RDF-17-01-05 (FK)

Start-up fund, Duke Kunshan University (FK)

Wang-Cai and Synear programs, Duke Kunshan University (FK)

Dachuang innovation program, Duke Kunshan University (FK)

## Author contributions

Conceptualization: GW, FK

Methodology: GW, MM, SK, PM,

Investigation: GW (conducted majority of experiments)

Visualization: GW

Supervision: FK

Writing—original draft: GW, FK

Writing—review & editing: GW, FK

## Data and materials availability

All data are available in the main text or the supplementary materials.

## Extended Data

**Extended Data Figure 1.**
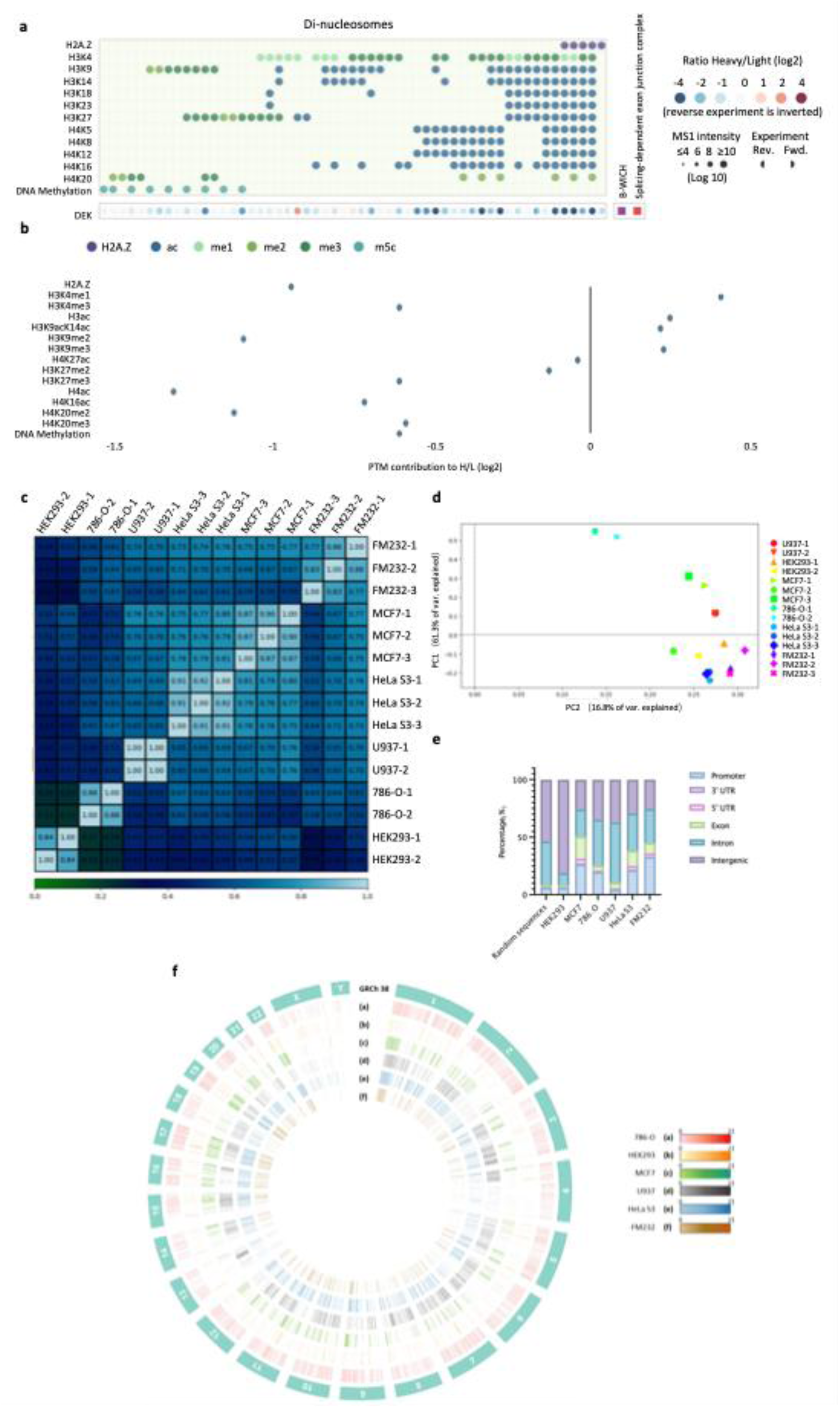
DEK binding preferences for modified nucleosomes and cross-cell-type comparison of DEK ChIP–seq occupancy. **a,** MARCS di-nucleosome library readout for DEK showing binding responses across nucleosomes bearing the indicated histone modifications and DNA methylation. Dot color indicates Heavy/Light ratio (log2) and dot size reflects MS1 intensity; forward and reverse experiments are shown (reverse experiment inverted). Adapted from the MARCS browser (Decoding chromatin states by proteomic profiling of nucleosome readers; https://marcs.helmholtz-munich.de/). **b,** Chromatin feature effect estimates for DEK derived from MARCS measurements, reported as PTM contributions to the Heavy/Light ratio (log2) for the indicated chromatin features. Adapted from the MARCS browser as in (a). **c,** Spearman correlation heatmap of genome-wide DEK ChIP–seq occupancy across datasets (U937, HEK293, MCF7, 786-O, HeLa S3, and FM232; replicates as indicated). **d,** Principal component analysis (PCA) of DEK ChIP–seq occupancy across the same datasets (PC1 and PC2 variance explained as shown). **e,** Genomic feature distribution of DEK ChIP–seq peaks across datasets shown as stacked bar plots (Promoter, 5′ UTR, 3′ UTR, Exon, Intron, Intergenic), with randomized genomic regions as background. **f,** Circos plot summarizing genome-wide DEK occupancy across datasets (GRCh38/hg38); color density indicates relative enrichment, with tracks corresponding to 786-O (a), HEK293 (b), MCF7 (c), U937 (d), HeLa S3 (e), and FM232 (f).

**Extended Data Figure 2.**
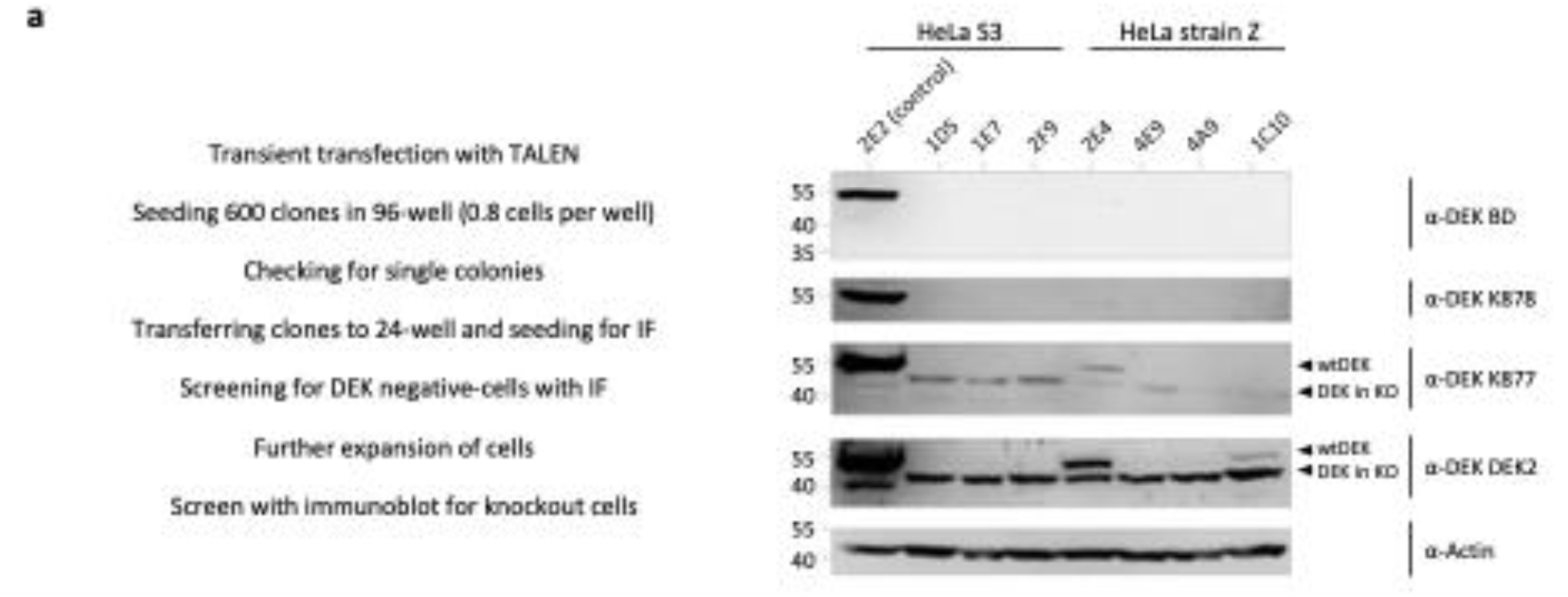
Generation and screening of the 1E7 HeLa S3 DEK-edited clone expressing truncated DEK. **a.** Transient TALEN transfection workflow used to derive clonal HeLa S3 cell lines: limited dilution (600 clones seeded in 96-well plates; ∼0.8 cells per well), identification of single colonies, expansion, and sequential screening by immunofluorescence (IF) for DEK-negative cells, followed by immunoblot validation. **b**, Immunoblots of candidate clones (including 2E2 control and 1E7) were probed with multiple DEK antibodies (α-DEK BD, α-DEK K878^22^, α-DEK K877^22^, and α-DEK2^73^) to assess DEK species, with actin as a loading control. All selected cell lines showed absence of any DEK-specific signal with the DEK BD-specific antibodies. However, two polyclonal antibodies (K877 and DEK2) detected a DEK-specific product in all lines. Interestingly, this band did not resemble an already described breakdown product of DEK^74^, which can be seen in the 2E2 control cell line at around 40 kDa. The fragment in the DEK knockout cells migrates faster than wildtype DEK, yet slower than the breakdown product. Transduction of cells with lentivirus delivering an shRNA targeting the 3’-end the *DEK* mRNA resulted in complete absence of this fragment (data not shown), suggesting expression of a N-terminally truncated DEK fragment, which is probably derived from an alternative transcriptional start site (ENCODE (https://genome.ucsc.edu/ENCODE/). Given that the intensity of this product is ∼ 10 % when compared to full length DEK, this may indicate that the fragment is expressed from one allele. Taken together, the TALEN approach indeed yielded cells lacking expression of full length DEK. However, a remaining DEK fragment of currently unknown nature was observed in all investigated cell lines.

**Extended Data Figure 3.**
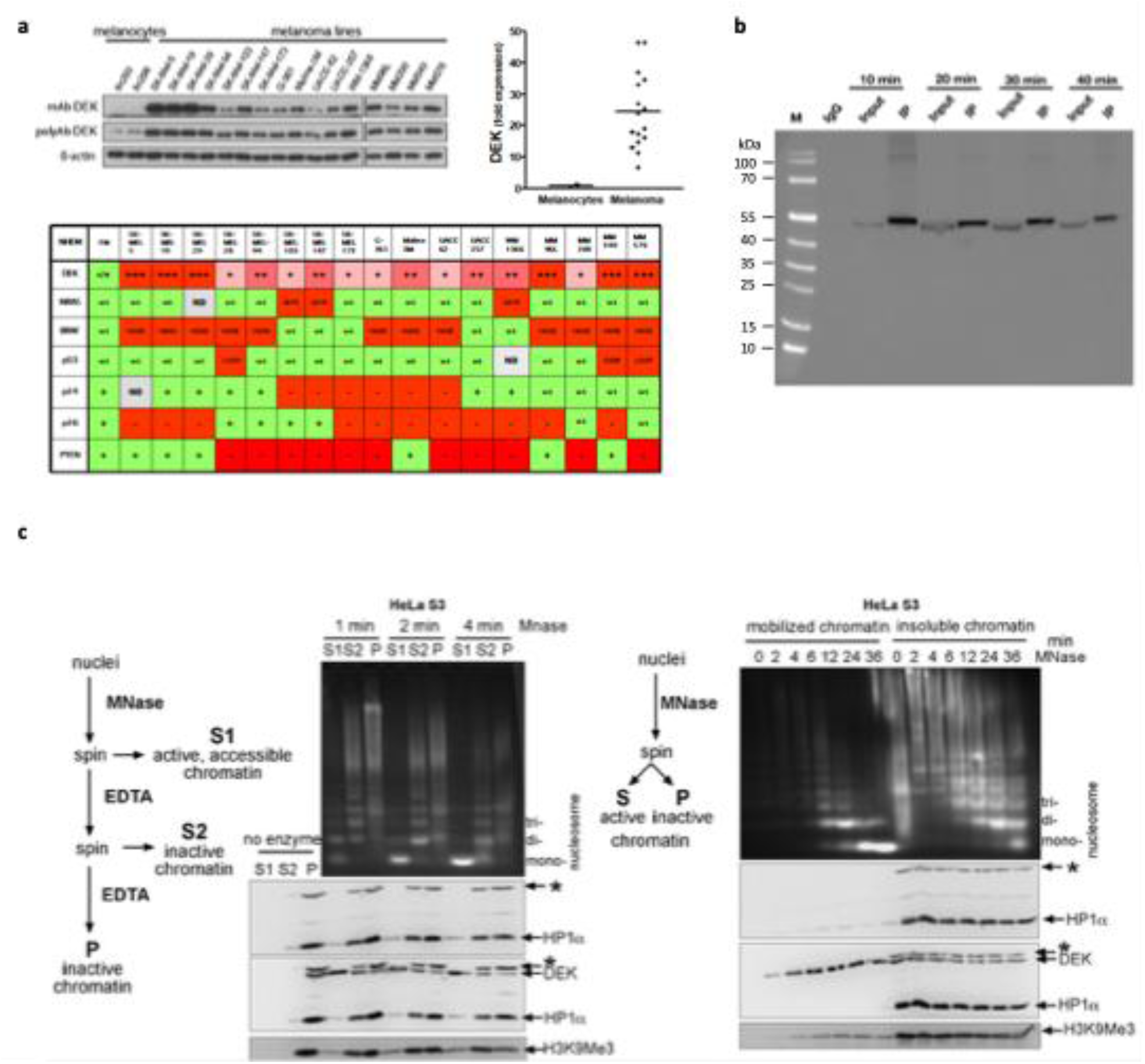
DEK abundance in melanoma models and technical controls for DEK chromatin assays. **a,** Immunoblot analysis of DEK protein levels in primary melanocytes and melanoma cell lines (including SK-Mel-19, SK-Mel-29, SK-Mel-103, UACC-62 and UACC-257) using monoclonal and polyclonal anti-DEK antibodies, with β-actin as loading control; the genetic background/driver alteration summary for the same panel of lines is shown as a heatmap. Figure partially adopted from^7^ **b,** Formaldehyde crosslinking time-course control for DEK ChIP. UACC-257 cells were crosslinked with 1% formaldehyde for the indicated times points, quenched, and processed for DEK ChIP; immunoprecipitated material was analyzed by SDS-PAGE and immunoblotting using DEK-specific antibodies. **c,** MNase accessibility-based chromatin fractionation schemes established in HeLa S3 cells. Left panel: a differential MNase-based extraction method as outlined in the graph on the left. Right panel: a time course MNase digestion regimen was applied as indicated on the left. Resulting samples were analyzed by agarose gel electrophoresis (top) and immunoblotting using DEK, HP1α, and H3K9me3-specific antibodies (* indicates a HP1α-antibody-specific cross reaction). Representative results of many repetitions are shown (see also^23^).

**Extended Data Figure 4.**
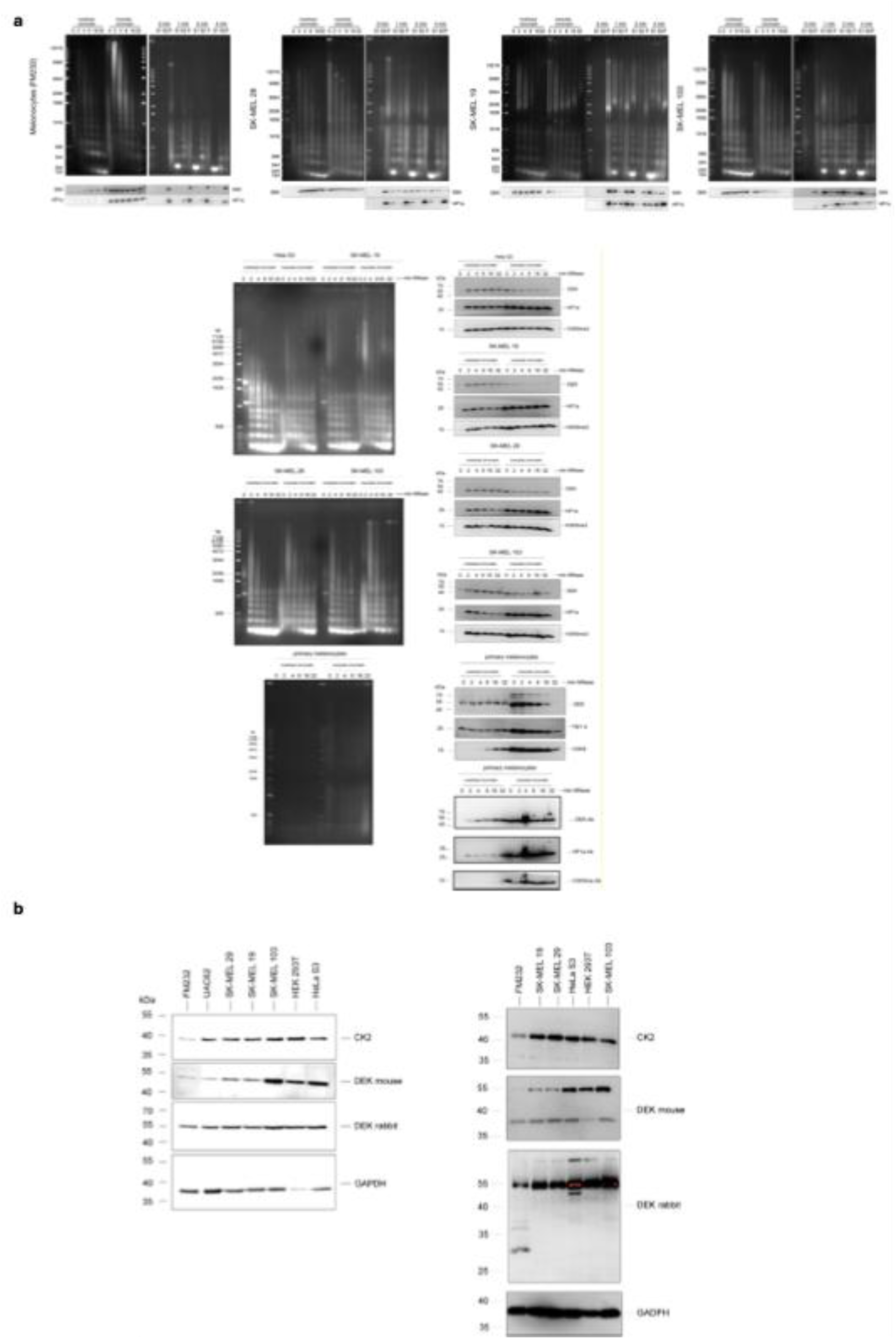
Chromatin fractionations and CK2 level in melanoma cell lines. **a**. MNase accessibility-based chromatin fractionation regimens were carried out and analyzed as detailed in Extended Data Figure 3c using the indicated cell lines or primary melanocytes. Representative results of many repetitions are shown. Please note that the degree of DEK distribution between accessible and inaccessible chromatin varies, yet the overall tendency is consistent. **b**. Immunoblot of total cell lysates from primary melanocytes and melanoma cell lines (as indicated) probed with a phosphorylation-preferring monoclonal DEK antibody and a polyclonal DEK antibody, together with CK2; GAPDH serves as a loading control. The phospho-DEK–preferential signal is increased in SK-Mel-103, coinciding with elevated CK2 level.

**Extended Data Figure 5.**
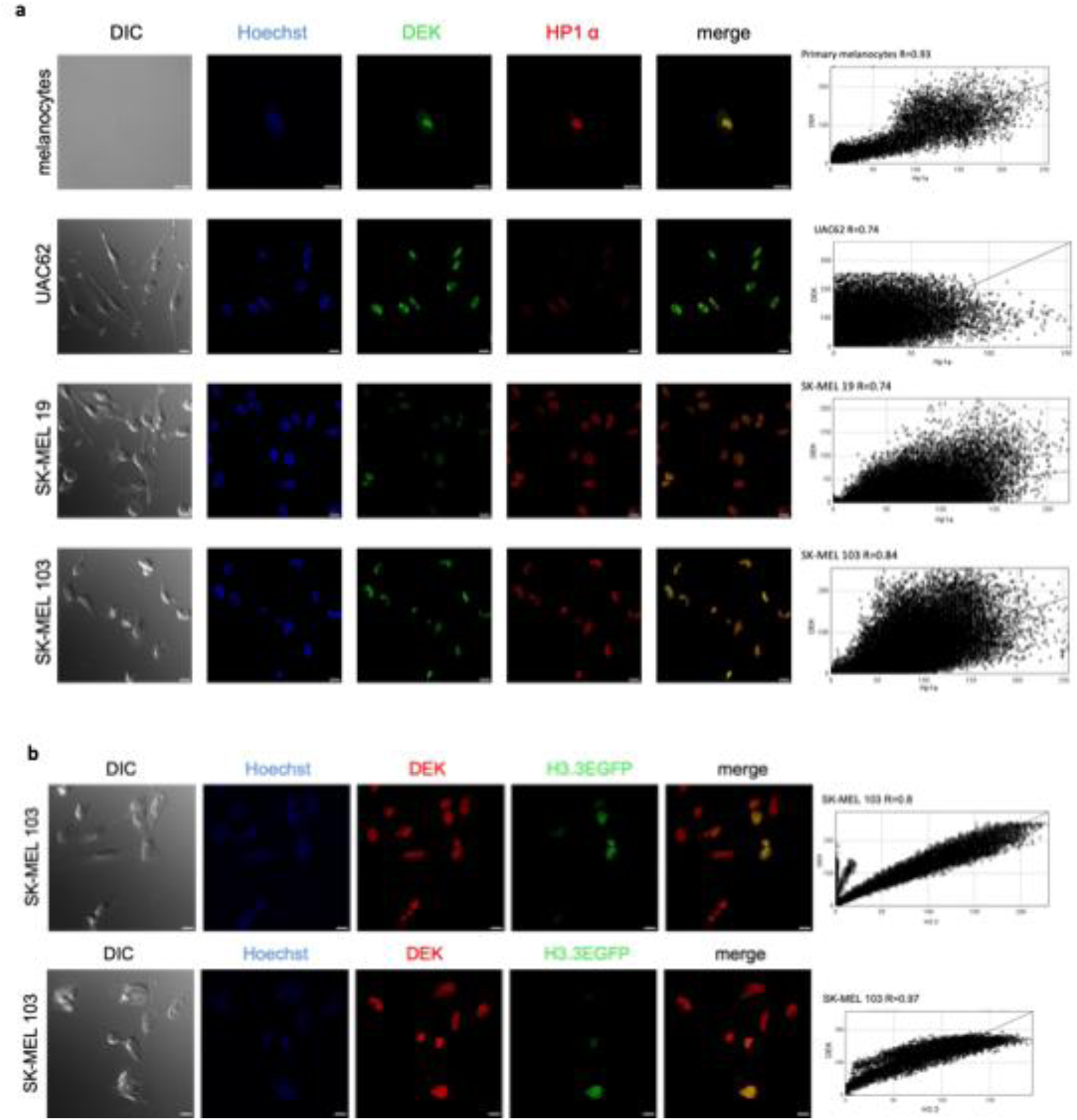
DEK differentially colocalizes with HP1α and Histone H3.3. **a**, Indicated cell lines were analyzed by immunofluorescence using indicated antibodies and correlations plots were performed using Correlation J in Image J. Shown is one representative replicate of at least 12 repetitions. **b**, SK-MEL 103 cells were transfected with the plasmid pH3.3GFP and subjected to immunofluorescence analysis and correlation analysis.

**Extended Data Figure 6.**
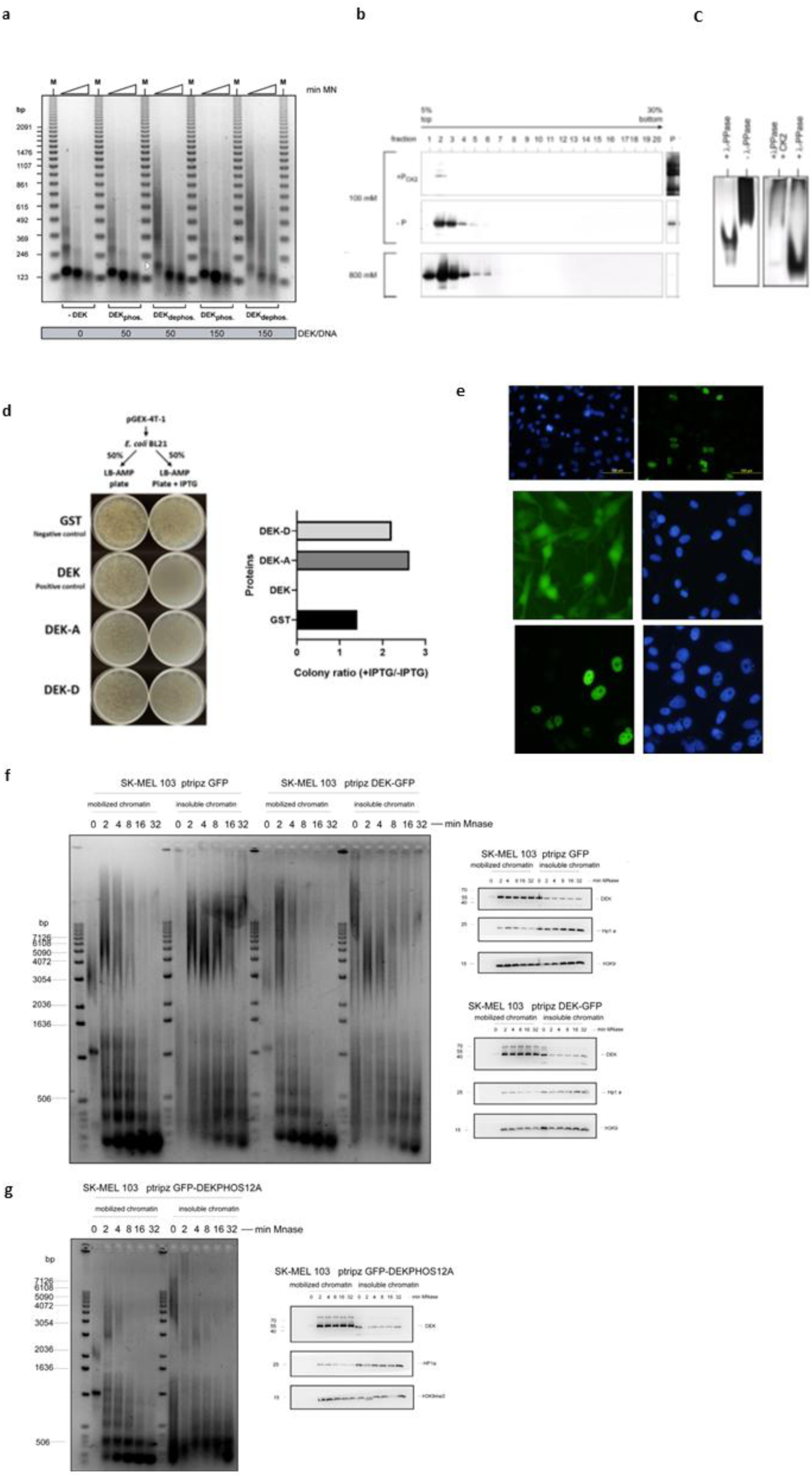
**a,** Micrococcal Nuclease Digestion of SV40 minichromosomes. Dephosphorylated and CK2-phosphorylated His-DEK were each incubated with 1 µg of salt-washed SV40 minichromosomes in nE100 for 1 h at 37°C. The individual samples were supplemented with 3 mM CaCl₂ and digested with 6 units of micrococcal nuclease for 1, 2, and 3 minutes. The reaction was stopped, the DNA was purified, separated on a 1.5% agarose gel, and visualized by SYBR Gold staining. **b**, Sucrose density gradients. wtHis-DEK was dephosphorylated using λ-phosphatase (-P) and *in vitro* phosphorylated by CK2 (+PCK2). Both samples were centrifuged on 5-30% sucrose gradients in nE100 (100 mM NaCl) for 14 hours at 36,000 rpm (SW-40) and 4°C. The gradients were fractionated into 20 fractions. The proteins were precipitated and analyzed by SDS-PAGE (10%) and immunoblotting with DEK-specific antibodies. The pellet fractions from the gradients were dissolved in SDS and analyzed as described above. Lower panel: sample was prepared as in (+PCK2) and sedimented on a 5-30% sucrose gradient in nE800 (800 mM NaCl). **c**, Native acrylamid gel electrophoresis. Dephosphorylated His-DEK (+ λ-PPase), untreated His-DEK (- λ-PPase), and CK2-phosphorylated His-DEK (+ λ-PPase, + CK2) were separated on a standard Laemmli polyacrylamide gel without SDS. Proteins were detected by silver staining. **d**, Bacterial Growth Inhibition Screen (BGIS) was carried out as described in ^19, 75^. pGEX-4T1 vectors containing full length DEK (DEK), S-287/288-A (DEK A), or S-287/288-D mutations were transformed into *E. coli* BL21 and plated on LB-AMP or LB-AMP-IPTG plates and incubated at 37C for 16 hours. Colony ratio between – and + IPTG was established using ImageJ. **e**, SK-MEL-103 cells were transduced with a pTRIPZ plasmid either expressing GFP or GFP-DEK and micrographs were taken 4 hours thereafter. **f**, SK-MEL-103 cells lentivirally transduced with pTRIPZ vectors expressing either GFP or DEK-GFP were selected, incubated with doxycycline for 24 hours and subjected to MNase time course digestions. Resulting DNA and proteins were analyzed by agarose gel electrophoresis and immunoblotting using indicated antibodies. **g**, SK-MEL-103 cells lentivirally transduced with a pTRIPZ vector expressing DEK-GFP S-11-A mutant: S-227, 230, 231, 232, 243, 244, 251, 301, 303, 306, 307-A were selected, incubated with doxycycline for 24 hours and subjected to MNase time course digestions and analyzed as in f.

**Extended Data Figure 7.**
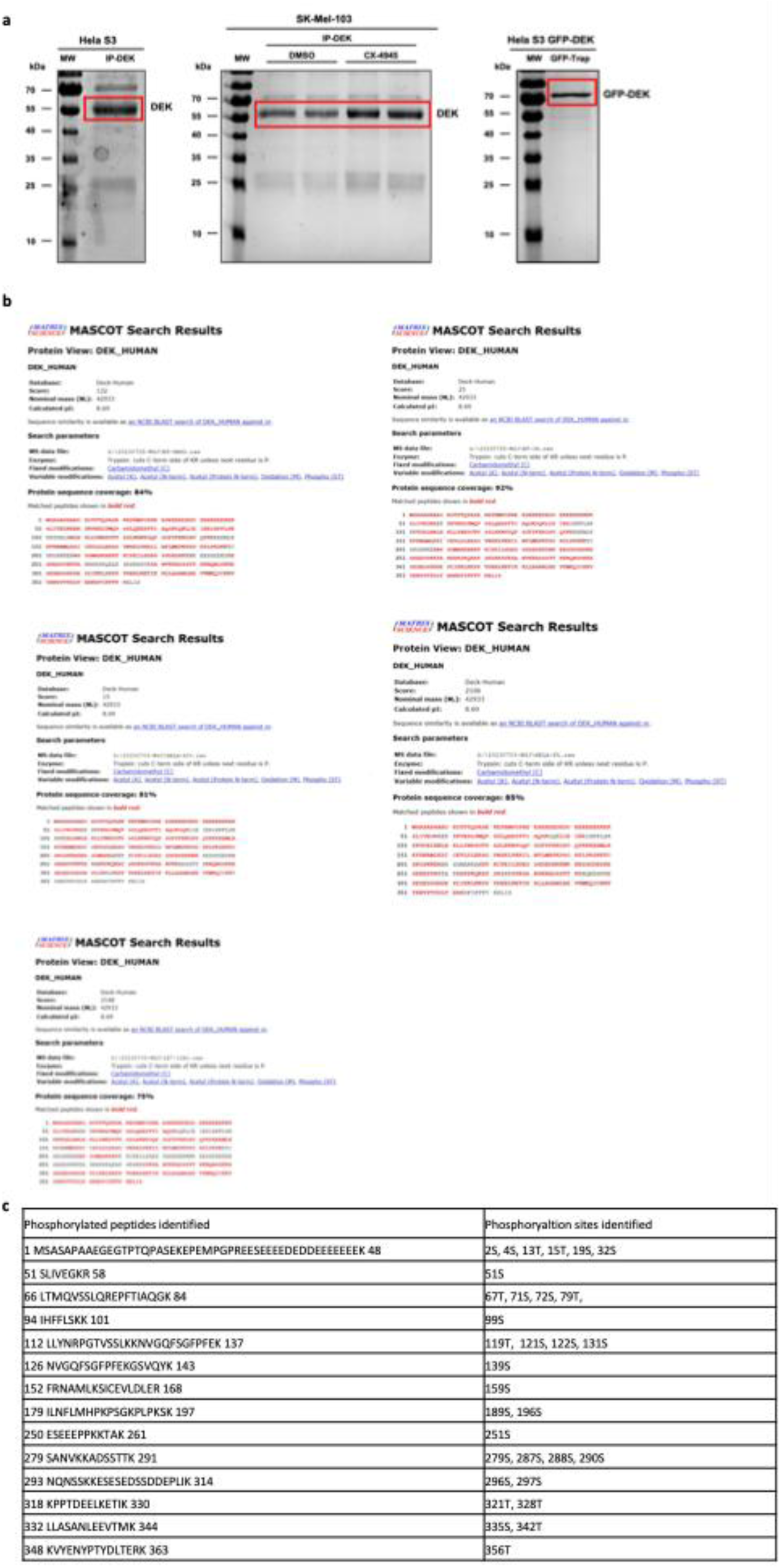
Mass spectrometry. **a,** Gels of immunoprecipitated DEK or GFP-DEK from HeLa S3 and SK-Mel-103 cells prepared for mass spectrometry analysis. Endogenous DEK or GFP-tagged DEK (GFP-DEK) was immunoprecipitated from HeLa S3 or SK-Mel-103 cells using DEK-specific antibodies or GFP-Trap beads, respectively. (**Left**) IP of endogenous DEK from HeLa S3 cells. (**Middle**) IP of DEK from SK-Mel-103 cells treated with DMSO or the CK2 inhibitor CX-4945. (**Right**) GFP-DEK immunoprecipitated from HeLa S3 cells expressing GFP-tagged DEK using GFP-Trap. All samples were resolved by SDS-PAGE and stained with Coomassie Brilliant Blue. The major DEK or GFP-DEK bands (highlight with red box) were excised for downstream mass spectrometry (MS) analysis to investigate phosphorylation sites. **b**, Mascot search results of DEK immunoprecipitates. Representative MASCOT search results from LC-MS/MS analysis of DEK protein treated with DMSO and CX-4945 from SK-Mel103, DEK and GFP-DEK from HeLa S3 cells, as well as GFP-DEK 12A mutant from 1E7. **c**, Phosphorylated DEK peptides detected in the indicated immunoprecipitates are shown, with modified residues and their positions in the DEK sequence annotated (S/T).

**Extended Data Figure 8.**
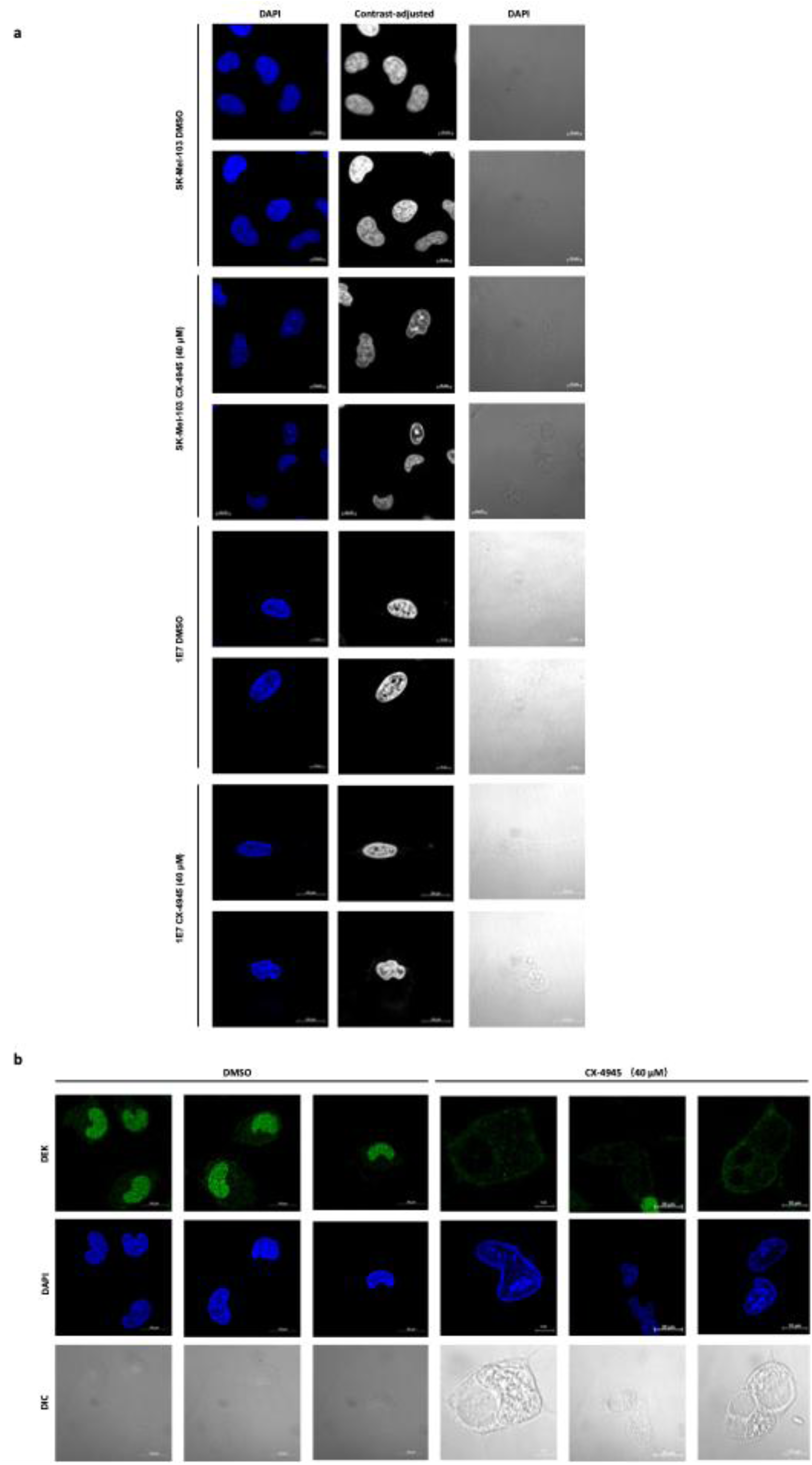
Additional confocal inmages supporting Fig. 7. **a,** Additional representative confocal images of nuclear chromatin morphology in SK-Mel-103 cells and DEK-depleted 1E7 cells following treatment with DMSO or the CK2 inhibitor CX-4945 (40 μM), corresponding to Fig. 7a. Shown are the DAPI channel, a contrast-adjusted grayscale rendering of the DAPI channel (for visualization of chromatin compaction), and DIC images. Scale bar, 10 μm. **b,** Additional representative immunofluorescence images of DEK staining in SK-Mel-103 cells following siRNA-mediated DEK knockdown and treatment with DMSO or CX-4945 (40 μM), corresponding to Fig. 7d. DAPI marks nuclei and DIC images are shown. Scale bar, 10 μm.

**Extended Data Figure 9.**
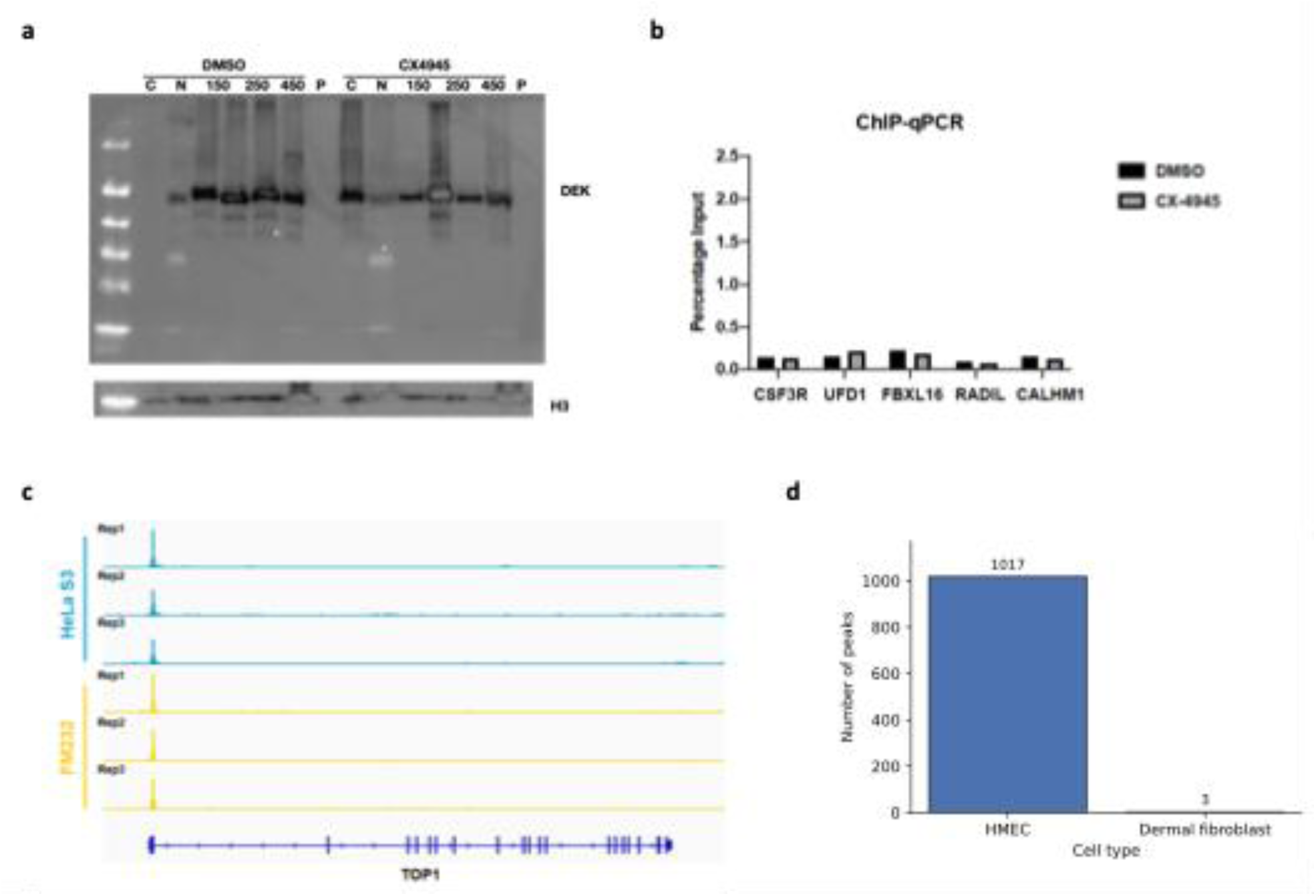
**a,** Replicate experiment relating to Fig. 5 d. Subcellular fractionation of SK-Mel-103 cells into cytosolic, nucleosolic and salt-extracted chromatin fractions (100, 250 and 450 mM NaCl), followed by immunoblotting for DEK. **b**, ChIP-qPCR validation was performed for CX-4945-specific DEK-accessible genes (UFD1, RADIL, FBXL16, CSF3R, and CALHM1) identified in SK-Mel-103 cells. ChIP was conducted in UACC-62 cells using antibodies against DEK and IgG as a negative control. This validation experiment was performed once to assess DEK enrichment at the corresponding loci**. c.** Representative genome browser tracks showing DEK ChIP**–**seq signal in HeLa S3 and FM232 cells at the TOP1 locus. **d**. Bar plot shows the number of DEK ChIP–seq peaks in human mammary epithelial cells (HMEC) and dermal fibroblasts.

